# Fecal microbial load is a major determinant of gut microbiome variation and a confounder for disease associations

**DOI:** 10.1101/2024.03.18.584290

**Authors:** Suguru Nishijima, Evelina Stankevic, Oliver Aasmets, Thomas S. B. Schmidt, Naoyoshi Nagata, Marisa Isabell Keller, Pamela Ferretti, Helene Bæk Juel, Anthony Fullam, Shahriyar Mahdi Robbani, Christian Schudoma, Johanne Kragh Hansen, Louise Aas Holm, Mads Israelsen, Robert Schierwagen, Nikolaj Torp, Manimozhiyan Arumugam, Flemming Bendtsen, Charlotte Brøns, Cilius Esmann Fonvig, Jens-Christian Holm, Trine Nielsen, Julie Steen Pedersen, Maja Sofie Thiele, Jonel Trebicka, Elin Org, Aleksander Krag, Torben Hansen, Michael Kuhn, Peer Bork, the GALAXY, MicrobLiver Consortia

## Abstract

The microbiota in individual habitats differ both in relative composition and absolute abundance. While sequencing approaches determine only the relative abundances of taxa and genes, experimental techniques for absolute abundance determination are rarely applied to large-scale microbiome studies. Here, we developed a machine learning approach to predict fecal microbial loads (microbial cells per gram) solely from relative abundance data. Applied to large-scale datasets (n = 34,539), we demonstrate that microbial load is the major determinant of gut microbiome variation and associated with numerous host factors. We found that for several diseases, the altered microbial load, not the disease itself, was the main driver of the gut microbiome changes. Adjusting for this effect substantially reduced the significance of more than half of the disease-associated species. Our analysis reveals that the fecal microbial load is a major confounder in microbiome studies, highlighting its importance for understanding microbiome variation in health and disease.

## Introduction

Shotgun metagenomic sequencing facilitates high-throughput profiling of complex microbial communities in environmental samples^1–3^. Applied to the human gut microbiome, metagenomics reveals its structure, function, and variations^4–6^, as well as its associations with host physiologies including diseases, immune function, and response to cancer therapy^7–11^. However, the microbial profile obtained from metagenomic analysis is inherently compositional, with the abundance of each microbial species represented in relative proportions (fraction of total reads)^12–14^. In such compositional data, changes in one microbial species result in concurrent relative changes in others, leading to negative correlation bias that can cause false positives and false negatives in association studies^12,13^. Moreover, sequencing data does not provide information on microbial load (i.e. the total absolute abundance of microbial species per gram or microbial density), which is closely associated with fecal transit time^15–17^, stool consistency^18^, water content^19^, and pH^20,21^ in the gut, and is a key ecological factor in shaping the diversity, metabolism, and inter-individual variation of the microbiome^19,22^.

To overcome these issues and to factor in total absolute abundances, various experimental methods are applied to microbiome studies, such as flow cytometry-based cell counting^19,23,24^, quantitative PCR^25–27^, or internal standard provision (e.g. spike-in DNA)^28–31^ that quantify the microbial load in environmental samples. Such additional data help avoid pitfalls associated with compositional data^13^ and link microbiome variation across individuals with changes in microbial load^19,32^. However, generating such quantitative profiles requires extra experiments that are labor-intensive, costly, and impractical for large-scale microbiome studies. Hence, the vast majority of public or ongoing metagenomic studies do not report associated microbial loads.

Here, we present a machine learning model capable of robustly predicting microbial load without requiring additional wet lab assays. Using large-scale paired datasets of metagenomes and microbial load data from two independent study populations (GALAXY/MicrobLiver and MetaCardis), we first train our model to predict the microbial load of a human fecal sample directly from relative microbiome profiles. We then demonstrate the utility of our model by applying it to a large-scale collection of public metagenomic datasets (n = 34,539), revealing novel associations between various host physiologies and microbial load. Furthermore, we show that microbial load is a major determinant of microbiome variation and frequently confounds disease associations of microbial species, with implications for biomarker development.

## Results

### Microbial loads are robustly predicted from the relative taxonomic and functional profiles of the microbiome

We based our analysis on fecal samples collected in two independent large-scale study populations by the GALAXY/MicrobLiver (n = 1,894, 46.7 ± 20.3 years old [mean ± s.d.], males 69.5%) and MetaCardis consortia (n = 1,812, 54.6 ± 13.0 years old [mean ± s.d.], males 44.8%)^33–35^. GALAXY/MicrobLiver encompassed various sub-cohorts including heterogeneous individuals such as healthy controls, early-to advanced-stage liver disease patients, individuals who participated in intervention trials, and children/adolescents with obesity (Methods, Supplementary Table 1). Meanwhile, MetaCardis focused on cardiometabolic disease patients (e.g. coronary artery disease, metabolic syndrome, type 2 diabetes, and severe/morbid obesity) as well as healthy individuals^33–35^ (Supplementary Figure 1 and Table 1). While the data on MetaCardis has been reported elsewhere^33–35^, we present here newly obtained metagenomes and flow cytometry-based cell count data from the GALAXY/MicrobLiver consortium (Supplementary Table 2). We obtained species-level taxonomic and functional (gene) profiles with a marker gene-based method using the mOTUs profiler^36^ and the Global Microbial Gene Catalog (GMGC)^37^, respectively. The microbial loads in the two study populations were significantly different (mean values were 6.5 ± 2.7e+10 and 11.1 ± 5.8e+10 for the GALAXY/MicrobLiver and MetaCardis study populations, respectively), suggesting possible study effects due to differences in experimental techniques used to measure load in respective study populations (Methods, Supplementary Figure 2). Nonetheless, taxonomic and functional profiles of the microbiome were consistently associated with the microbial loads in both study populations (Supplementary Figure 3, Supplementary Tables 3 and 4).

We first associated the experimentally measured microbial loads with three enterotypes^38,39^ (Methods). The microbial load was the highest in *Firmicutes* (*Ruminococcus*) enterotype followed by *Prevotella*, and *Bacteroides* enterotypes in both study populations (Figure 1A, B). Diversity indexes (e.g. Shannon diversity, species richness, and Simpson diversity) of the microbiome had consistent positive correlations with microbial load, with Shannon diversity showing one of the strongest positive associations in both study populations (Figure 1C). We next studied correlations between relative species abundance and total microbial load, and observed positive correlations for various uncultured species in *Firmicutes* phylum as well as short-chain fatty acid producers^40^ and slow-growing^41^ species (e.g. *Oscillibacter*, *Faecalibacterium,* and *Eubacterium* spp.). In contrast, we observed negative correlations for disease-associated species such as *Ruminococcus gnavus* (inflammatory bowel disease)^42,43^ and *Flavonifractor plautii* (colorectal cancer)^44^ (Figure 1C, Supplementary Table 3). Typical oral species also found in stool such as *Streptococcus* and *Veillonella* spp. were also negatively associated with the microbial load (Supplementary Figure 4).

**Figure 1.**
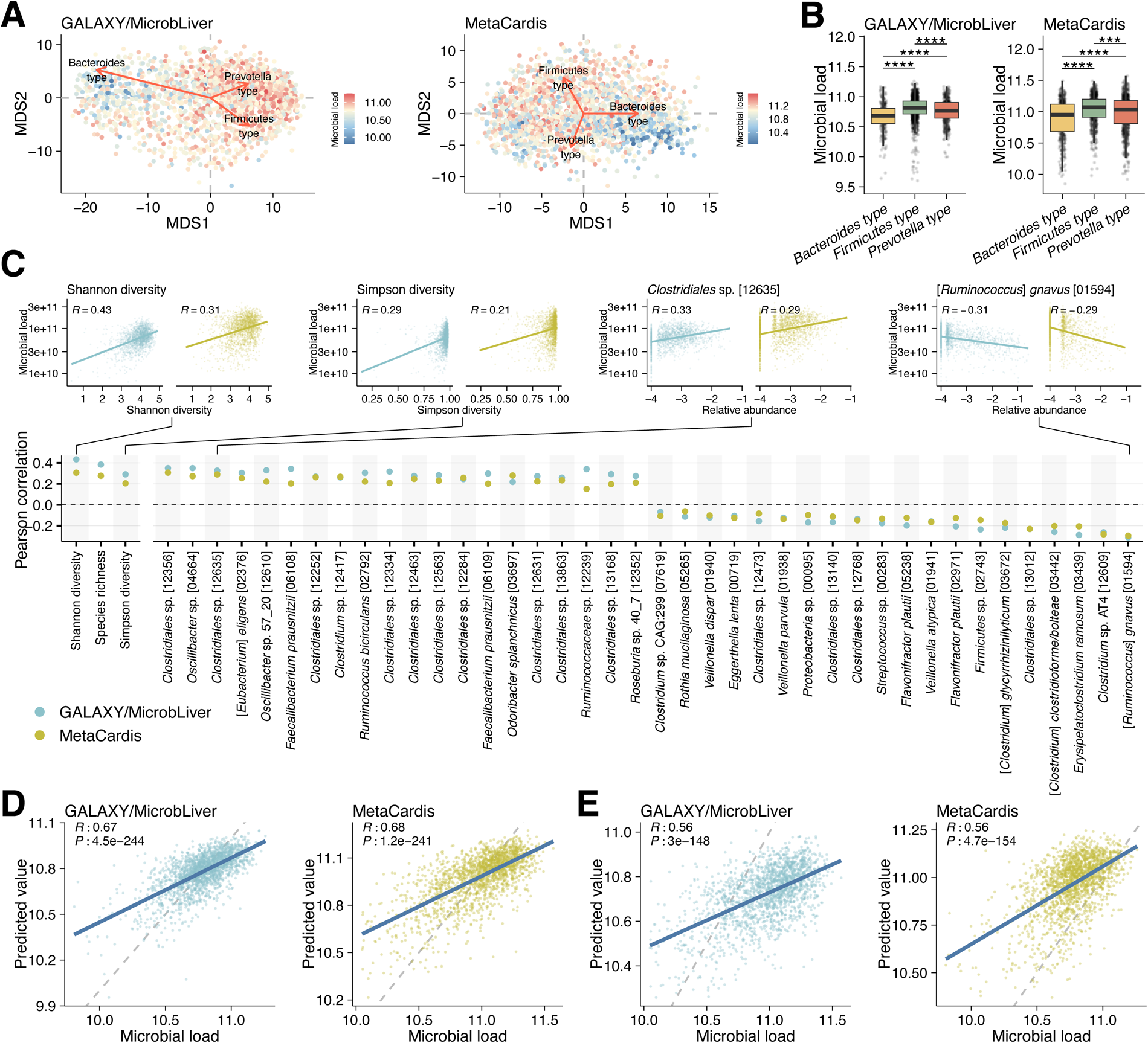
Machine-learning models robustly predict microbial loads of fecal samples. **A**, Multidimensional scaling plot of the species-level taxonomic profile of the microbiomes in the GALAXY/MicrobLiver (n = 1,894) and MetaCardis (n = 1,812) study populations. Each point represents a sample and the color shows the log10 transformed microbial load of the sample. Arrows represent the three enterotypes plotted by the envfit function in R. The direction of the arrow indicates the centroid of each enterotype, and the length indicates the strength of the correlation with the enterotype. **B**, Associations between the microbial loads and the enterotypes. Boxplots show the log10-transformed microbial load across the three enterotypes in each cohort. **** *P* < 0.0001, *** *P* < 0.001 (Wilcoxon rank-sum test). **C**, Pearson correlations between microbial load and relative abundances of microbial species (both values were log10 transformed). The three diversity indexes and the top 40 species with the highest correlations are shown. Scatter plots for the two diversity indexes and two microbial species are shown above the heatmap, as examples. **D**, **E**, Internal (**D**) and external (**E**) validation of the XGBoost prediction models. Scatter plots show the actual microbial load of fecal samples (log10) on the x-axis and predicted values on the y-axis. The solid blue lines show regression lines and the gray dashed lines represent 1:1 reference lines. For internal validation, the models were evaluated with a 5-times repeated 10-fold cross-validation. For external validation, the GALAXY/MicrobLiver and MetaCardis models were applied to each other’s datasets. Pearson correlation was used to evaluate the model’s accuracy.

When correlating the microbial loads with the relative functional profiles of the human gut metagenome, we found that microbial genes for lipopolysaccharide (LPS) biosynthesis were enriched in samples with low microbial loads in both study populations (Supplementary Figure 5, Supplementary Table 4). Similarly, genes for sugar metabolism including the phosphotransferase system, fructose/mannose metabolism, and glycan degradation were consistently associated with lower microbial loads in both study populations. On the other hand, genes involved in flagella assembly and bacterial chemotaxis were positively correlated with the high microbial load in both study populations (Supplementary Figure 5). As increased LPS levels in the gut could cause inflammation and diarrhea (i.e. shorter transit time)^45^, these genes might be also associated with fecal transit time.

As we observed strong associations between microbial load and relative gut microbiome profiles, we hypothesized that the microbial load of a fecal sample could be predicted from relative abundances of taxa. We thus trained eXtreme Gradient Boosting (XGBoost) regression models^46^ based on relative abundance of each microbial species as well as the Shannon diversity index (Methods). Internal 5-times repeated 10-fold cross-validation in each study population showed that both models predicted the microbial load with Pearson correlation coefficients of 0.67 ± 0.0068 and 0.68 ± 0.0069 (mean ± s.d.) for the GALAXY/MicrobLiver and MetaCardis study populations, respectively (Figure 1D). To evaluate the robustness of the model in an external dataset, we applied each model to the other dataset (Supplementary Figure 1) and found that both models again predicted the microbial loads significantly (Figure 1E, Pearson correlations = 0.56 for both the GALAXY/MicrobLiver and MetaCardis models). Functional profiles of the gut microbiomes also predicted the microbial loads with comparable accuracies to those trained by the species-level taxonomic profiles (Supplementary Figure 6A and B). These results demonstrated robust prediction of microbial loads in fecal samples from relative microbiome profiles obtained by metagenomic sequencing.

Since the GALAXY/MicrobLiver and MetaCardis study populations included individuals with different phenotypes and demographic factors (e.g. healthy adults, diseased patients, and children/adolescents, Supplementary Table 1), we next examined how prediction accuracy differed among these groups. Notably, we found that both models robustly predicted microbial load not only in healthy samples, but also in diseased samples that were not included in the model’s training (Supplementary Figure 7). Specifically, the MetaCardis model, which was trained on samples from healthy adults and cardiometabolic disease patients, showed comparative accuracy among the sub-cohorts in the GALAXY/MicrobLiver study population, such as the cohort of healthy individuals (GALA-ALD: Pearson correlation = 0.52), liver disease patients (GALA-ALD: 0.47, GALA-RIF: 0.58, and TIPS: 0.62), and children/adolescents (HOLBAEK: 0.53). Similarly, the GALAXY/MicrobLiver model also showed comparative accuracies for individuals with various diseases in the MetaCardis dataset, such as healthy individuals (Pearson correlation = 0.44), patients with coronary artery disease (0.43), diabetes (0.56), metabolic syndrome (0.48), and severe obesity/morbid obesity (0.63). As such, the models robustly predict microbial load even for samples with phenotypes not included in the training data.

To further explore the applicability to different sequencing technologies, we collected additional paired data of 16S rRNA gene sequencing and fecal microbial loads from two previous studies^19,24^ (Supplementary Table 5). The internal and external validations of the model between the two studies also demonstrated robust prediction of microbial load (Pearson correlations = 0.79 for the internal validation and 0.60 for the external validation, Supplementary Figure 6C and D), indicating that with sufficient data, fecal microbial loads can be predicted from different relative abundance measures.

### Predicted microbial loads are associated with various host factors

To discover associations between predicted microbial loads and host factors such as disease status, medication, and lifestyle, we collected public gut metagenomes from 159 previous studies across 45 countries (n = 27,832, 46.3 ± 19.3 years old [mean ± s.d.], 52.9% males, Supplementary Tables 6 and 7). Additionally, we collected metagenomes from two large population studies^47,48^: Japanese 4D cohort (n = 4,198, 66.4 ± 12.6 years old [mean ± s.d.], 58.8% males) and Estonian Microbiome cohort (n = 2,509, 50.0 ± 14.9 years old [mean ± s.d.], 29.7% males), in which deep phenotyping was performed (Supplementary Tables 6 and 8). Since the former data were derived from various smaller studies with less host intrinsic and extrinsic factor information, they were combined into a global dataset. We prepared species-level taxonomic profiles of each sample and predicted the microbial load using the MetaCardis prediction model (Methods, Supplementary Figure 1).

In the global dataset, samples from high-income countries showed significantly higher predicted microbial loads than those from low-income countries (Figures 2A and B). This difference could not be attributed to the potential bias that the model was trained on samples from high-income countries (Supplementary Figure 8), suggesting that factors associated with increased income such as lifestyle, diet, or hygiene affect the microbial load. In the Japanese 4D and Estonian Microbiome cohorts, the medication category showed the strongest association with the predicted microbial load among the metadata categories (Figure 2C, Supplementary Table 8), which was consistent with the strongest impact of medication on the relative gut microbiome profile observed previously^47^. Anthropometric factors and disease status followed in the Japanese 4D cohort, while diet and other factors ranked next in the Estonian Microbiome cohort. Among the available host factors in the Japanese 4D and Estonian Microbiome cohorts, 65 (26.3%) and 8 (3.2%) showed significant associations with the predicted microbial loads, respectively (FDR < 0.05, Figure 2D, Supplementary Table 9). In the three datasets, the self-reported Bristol stool scale (an index that classifies categories of fecal consistency) showed consistent negative correlations with the predicted microbial loads (Figure 2E, Supplementary Figure 9A). The frequency of defecation, surveyed in the Estonian Microbiome cohort, was also negatively associated with the microbial load (Figure 2F). These results are consistent with findings from previous studies suggesting that fast transit time (e.g. frequent defecation and diarrhea) reduces microbial loads while slow transit time (e.g. infrequent defecation and constipation) increases microbial loads since fecal bacteria grow along the gastrointestinal tract gradually^18,22^. Age was positively associated with the microbial load in both Japanese 4D and global datasets, but not in the Estonian Microbiome dataset (Supplementary Figure 9B). Overall, elderly individuals (>70 years old) had 9.7% higher microbial load than younger individuals (<30 years old) in the combined datasets (Figure 2G). Sex was consistently associated with the microbial load in all three datasets (Figure 2H, Supplementary Figure 9C), with women having a 3.5% higher microbial load than men on average. These results are consistent with epidemiological studies that showed slower transit time in elderly people and females^49,50^. Interestingly, elderly individuals and females showed higher microbiome diversity than younger individuals and males as observed in previous studies^51,52^, while the strength of these associations decreased once adjusted for the effect of the microbial load (Supplementary Figure 10). This suggests that the higher microbial load or slower transit time contributes to increased gut microbiome diversity in elderly individuals and females. Other factors significantly associated with microbial load in the Japanese 4D cohort included various medications (e.g. platelet aggregation inhibitors, aminosalicylic acid, osmotic laxatives), diseases (e.g. Crohn’s disease, ulcerative colitis, HIV infection), diet (e.g. fruits, mushrooms, green tea, vinegar), and lifestyle (e.g. alcohol consumption) (Figure 2D).

**Figure 2.**
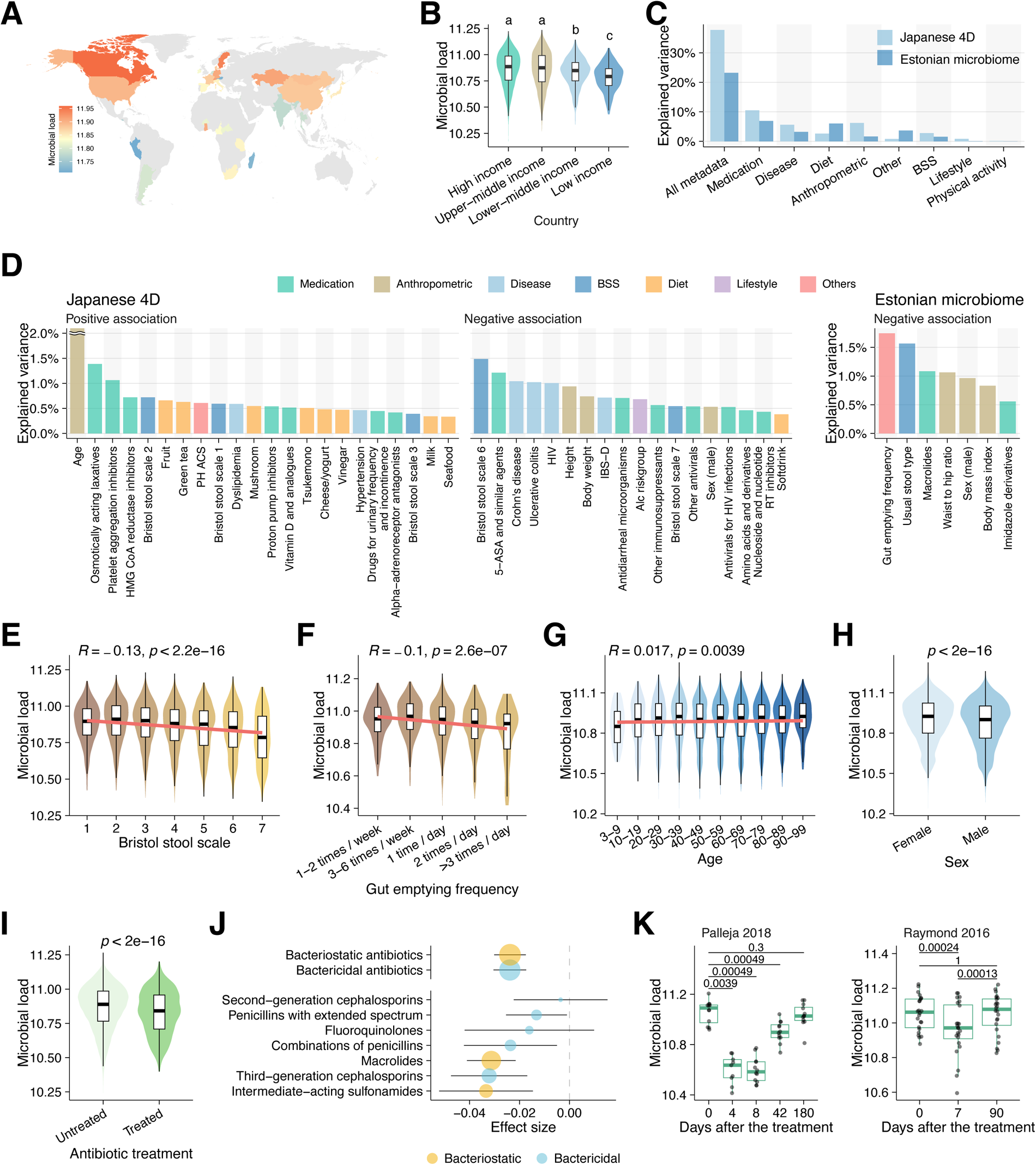
Predicted microbial loads are associated with various host factors. **A**, Predicted microbial loads of the collected metagenomes across different countries. Individuals treated with antibiotics and those with any diseases were excluded. The average microbial loads of the 34 countries with at least 20 individuals are shown. **B**, Comparison of the predicted microbial loads among four groups of countries divided by economic size. Definitions of the groups were obtained from the World Bank. The letters above the boxes (a, b, and c) indicate statistically significant differences (*P* < 0.01) between groups with different letters. **C**, Associations between the predicted microbial load and various host factors in the Japanese 4D and Estonian Microbiome cohorts. The explained variances by the host factors (coefficient of determination) were assessed by linear regression models including these host factors as explanatory variables and the log10 transformed microbial load as a response variable. **D**, Associations between the predicted microbial load and each host factor. The explained variance was assessed by linear regression models and the top 40 factors with the strongest associations in the Japanese 4D cohort (FDR < 0.05) and eight factors with FDR < 0.05 in the Estonian Microbiome cohort are shown in the figure. For visualization, the explained variance for age in the Japanese 4D cohort (2.8%) is plotted above 2.0% on the y-axis. **E**, **F**, **G**, **H, I**, Correlations between the predicted microbial load and various host factors, such as Bristol stool scale (**E**), gut emptying frequency (**F**), age (**G**), sex (**H**), and antibiotics (**I**). Associations were evaluated with Pearson correlation for **e**, **f**, **g,** and Wilcoxon rank-sum test for **H** and **I**. **J**, Effects of different types of antibiotics on the predicted microbial load in the Japanese 4D, and Estonian Microbiome cohorts. Each circle shows the effect size (beta coefficient) determined by a linear regression analysis and the error bar represents 95% confidence intervals. Blue and yellow colors show bactericidal and bacteriostatic antibiotics, respectively. **K**, Recovery of the predicted microbial load after antibiotic treatment. Boxplot showing the predicted microbial load of each individual at the respective time point. The datasets were collected from Palleja *et al.* (n = 12) and Raymond *et al.* (n = 24) studies. Numbers in the plot indicate the *P* values of each time point in comparison with the baseline (paired Wilcoxon rank-sum test).

Among medications, antibiotics substantially disrupt the microbial community in the human gut^53,54^, but only a few small-scale studies quantified changes in the microbial load^32,55^. As expected, recent antibiotic treatment was negatively associated with predicted microbial loads in all three datasets (Figure 2I and Supplementary Figure 9D). Using detailed information on classes of antibiotics from the Japanese 4D and Estonian Microbiome cohorts, we found that many had significant impact on the microbial loads, such as sulfonamides, third-generation cephalosporins, macrolides, and fluoroquinolone (Figure 2J). We did not find any differences between bactericidal (i.e. those that kill bacteria) and bacteriostatic (i.e. those that prevent bacterial growth) antibiotics, in line with recent findings that there might not be such a clear separation between bactericidal and bacteriostatic groups^56^. To further explore changes in the microbial loads, we focused on two public time-series metagenomic datasets, with data up to 180 days post-antibiotic treatment^54,53^. In one of these^54^, individuals were treated with a combination of three broad-spectrum antibiotics (vancomycin, gentamicin, and meropenem) while in the other^53^, individuals were treated with a second-generation cephalosporin (cefprozil). We found that the microbial loads gradually recovered after the treatment and returned to the baseline level in 180 and 90 days, respectively (Figure 2K). However, the microbial load was still significantly reduced at day 42 following combinatorial treatment^54^. These results suggest that recovery of microbial load after antibiotic treatments takes at least several weeks. This is consistent with studies on relative abundances, reporting recovery only after months^54,57^.

### Numerous diseases are associated with altered microbial loads

Identification of disease-associated gut species is an important step in developing microbial biomarkers, investigating the etiology of diseases, and developing targeted therapies^11,58,59^. However, the association between microbial load and disease is still largely unexplored, except in a few cases where microbial loads were experimentally determined^19,32^. To evaluate associations between various diseases and the fecal microbial loads, we performed a large-scale case-control analysis by combining the datasets from the global and Japanese 4D datasets, capturing sufficient data for 26 diseases (i.e. >50 cases and controls for each disease) across 11,807 cases and 17,118 controls (Supplementary Table 10, Supplementary Figure 11). The analysis revealed that the majority of diseases (14/26) were significantly associated with microbial load (FDR < 0.05). Nine of the significantly associated diseases showed negative associations with microbial load while five showed positive associations (Figure 3, Supplementary Table 11). The negatively-associated diseases included Crohn’s disease, ulcerative colitis, liver cirrhosis, *C. difficile* infection, and HIV infection, all of which are frequently associated with diarrhea^60–62^. On average, these patients with these conditions had 17.3% lower microbial loads than controls (Supplementary Table 11). Positively associated diseases included slow transit constipation, along with conditions often associated with constipation such as multiple sclerosis, colorectal cancer, and hypertension^63,64^. On average, these patients exhibited 7.7% higher microbial load than the controls. In irritable bowel syndrome (IBS), which is classified into different subtypes according to symptoms, the diarrhea type (IBS-D) showed a significant negative correlation (FDR = 3.6e-07), while the constipation type (IBS-C) showed a significant positive correlation with microbial load as expected (FDR = 0.042, Figure 3).

**Figure 3.**
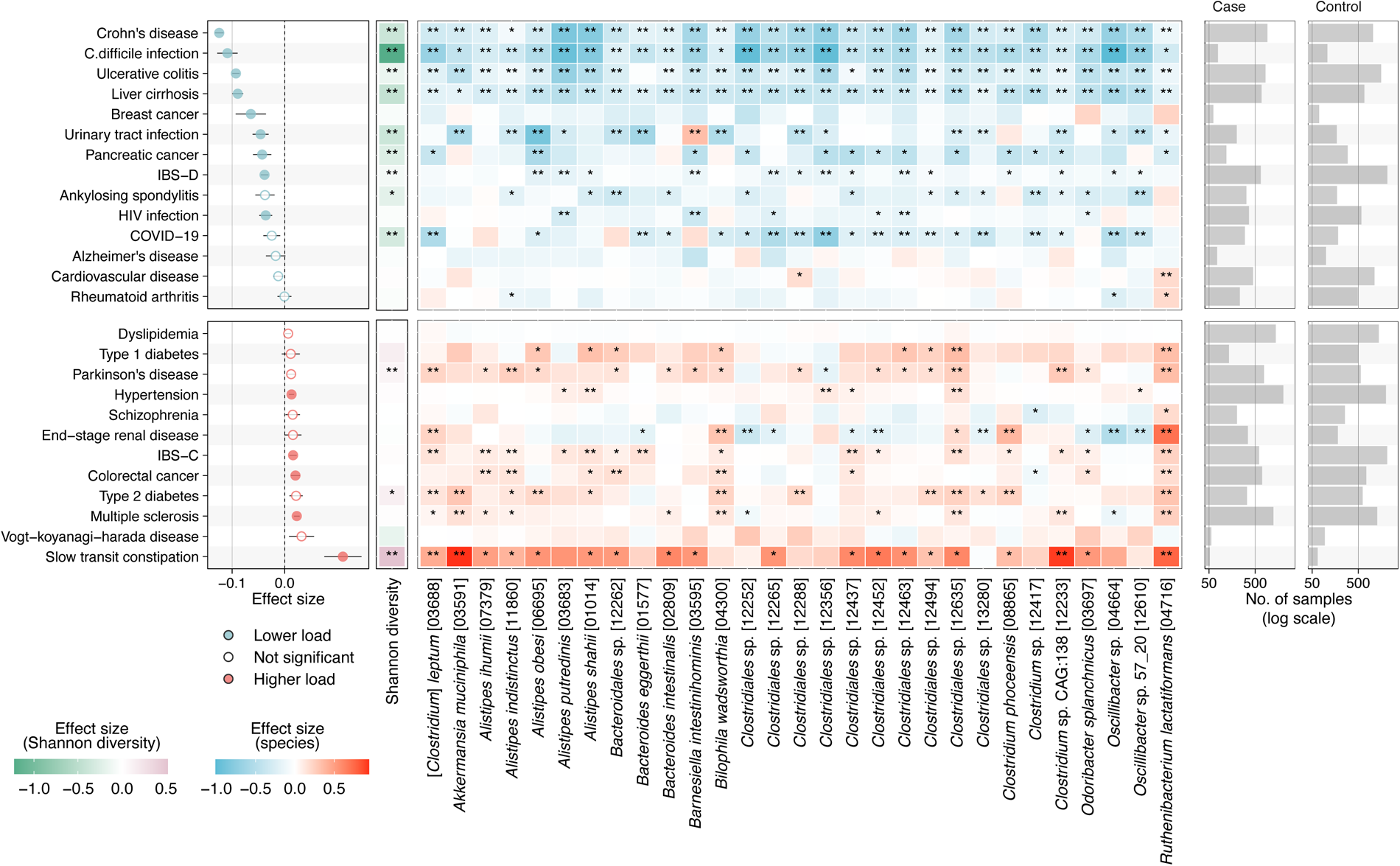
Predicted microbial loads are associated with various diseases. The left forest plot shows the effect sizes of the disease on the predicted microbial load. Blue and red colors represent negative and positive associations with the microbial load compared to the controls, respectively. Filled and empty circles represent significant (FDR < 0.05) and non-significant (FDR > 0.05) diseases, respectively. Effect sizes were assessed by a linear regression model including log10 transformed microbial load as a response variable and disease status (i.e. case or control) and each study as explanatory variables. The middle heatmap shows enrichments and depletions of microbial species across different diseases. Blue and red colors represent negative and positive associations, respectively, compared with the controls in each disease dataset. The top 30 species with the strongest differences (FDR < 0.05) in their effect sizes between positively-and negatively-associated diseases are shown. ** *P* < 0.01, * *P* < 0.05 (linear regression analysis). The right bar plot represents the number of samples included in the comparison.

To further characterize microbiome profiles in these diseases, we performed a meta-analysis of the relative microbial compositions between cases and controls, and defined microbial signatures for each disease based on the coefficient for each species obtained from the regression model (Methods). Comparison of the signatures between positively and negatively-associated diseases revealed a significant difference between the two groups (*P* = 0.0002, Supplementary Figure 12). The majority of the negatively-associated diseases was characterized by significantly less diverse microbiomes (e.g. in Crohn’s disease, *C. difficile* infection, and ulcerative colitis), which was in line with previous findings that diseases accompanied by diarrhea commonly present reduced microbiome diversity^65^. In contrast, some of the positively-associated diseases showed significantly increased microbiome diversity (e.g. slow transit constipation, type 2 diabetes, and Parkinson’s disease). Additionally, we identified 87 species that distinguished positively- and negatively-associated diseases (FDR < 0.05), such as *Alistipes (A. putredinis, A. indistinctus and A. shahii)*, *Bacteroides (B. eggerthii, B. intestinalis, and B. clarus)*, and *Eubacterium* spp. (*E. siraeum* and *E.* sp. CAG:202) as well as uncultured *Clostridiales*. The majority of these species were consistently depleted in the patients with negatively-associated diseases, while enriched in those with positively-associated diseases (Figure 3, Supplementary Table 12). These species also included *Bilophila wadsworthia*, a hydrogen sulfide-producing bacteria that may cause systemic inflammation^66,67^, and *Akkermansia muciniphila*, a potential beneficial microbe that may enhance the gut barrier integrity^68^. An unclassified *Burkholderiales* species was the only species consistently enriched in the negatively-associated disease patients while depleted in positively-associated disease patients (Supplementary Table 12). The presence of these consistent disease-microbe associations across different diseases suggests that they are confounded by the microbial load.

### Microbial load substantially confounds disease-microbe associations

To disentangle species association with disease from those with microbial load in the case-control analyses, we next incorporated microbial load as a covariate in a regression model, which is a method to effectively adjust for such confounding effects in microbiome studies^69,70^ (Methods). We excluded Vogt-Koyanagi-Harada disease and Alzheimer’s disease from the following analyses since no significant species were identified in these two diseases (FDR > 0.05). The adjustment led to a considerable reduction in the effect size of the disease-species associations and their statistical significance (in terms of p-value) in several diseases. This was especially the case for seven diseases, namely Crohn’s disease, ulcerative colitis, liver cirrhosis, IBS-D, breast cancer, *C. difficile* infection, and slow transit constipation (Figure 4). For these conditions, the adjustment led to a decrease in the average effect size on species by 21.9 to 49.9% (35.5% on average, Figure 4A), and consequently, 23.6 to 75.0% (48.0% on average) of the previously significant disease-species associations (FDR < 0.05) were no longer significant (FDR > 0.05, Figure 4B and C). Of these seven diseases that were particularly affected by the adjustment, six, except for slow transit constipation, were the ones negatively associated with microbial load. On the other hand, several diseases positively associated with microbial load, such as end−stage renal disease, colorectal cancer, and multiple sclerosis showed slight increases in the number of significantly associated species with them (Figure 4C).

**Figure 4.**
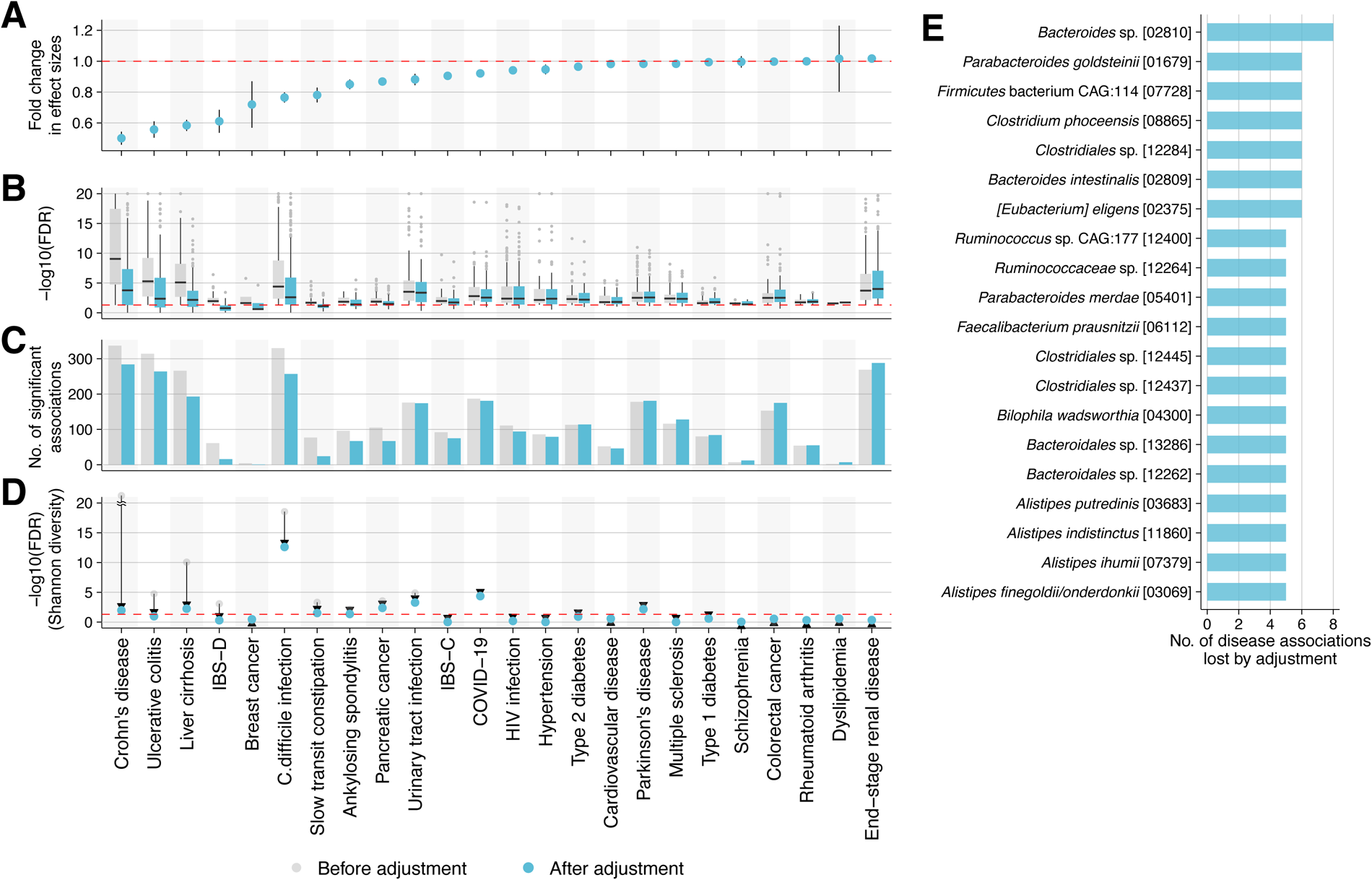
Microbial loads confound disease-microbe associations. **A,** Fold change in effect size before and after adjustment for species that were significantly associated with the disease (FDR < 0.05) before the adjustment for the microbial load. The y-axis shows the geometric mean of the ratio of the effect size on the species before and after the adjustment. The error bars show the 95% confidence interval of the geometric mean. Associations between the disease and species abundances were assessed by linear regression analysis with and without the microbial load as a covariate (Methods). Results for 24 diseases are shown in the plot as Vogt-Koyanagi-Harada disease and Alzheimer’s disease had no significant associations with any species (FDR > 0.05). **B,** Comparison of the statistical significance (i.e. FDR) of species before and after the adjustment. For visualization, the maximum on the y-axis was set at 20 (i.e. FDR = 1e-20), and some extremely lower FDRs were plotted there. **C**, Comparison of the number of significantly associated species (FDR < 0.05) before and after the adjustment. **D**, Comparison of the statistical significance of the Shannon diversity before and after the adjustment. Arrows represent the changes in the FDR before and after the adjustment. Red horizontal line represents FDR = 0.05. For visualization, the FDR for Crohn’s disease before adjustment (2.2e-25) is plotted above 20 on the y-axis. **e**, The top species (n = 20) that lost their significant associations to at least 5 of the 26 diseases due to the adjustment.

Microbial species that lost their significance across different diseases after the adjustment included *Clostridium phoceensis, Bacteroides intestinalis, Eubacterium eligens, Parabacteroides merdae, and Faecalibacterium prausnitzii* (Figure 4E, Supplementary Figure 13, and Supplementary Table 13), suggesting that significant changes in their abundances are mainly due to the changes in the microbial load rather than the disease status. The majority of these species substantially affected by the adjustment were those depleted in the disease patients. In contrast, several species significantly enriched in disease patients were not substantially affected by the adjustment, such as *Fusobacterium nucleatum* in colorectal cancer, *Flavonifractor plautii* in Crohn’s disease and ulcerative colitis, and *Streptococcus anginosus* in liver cirrhosis and pancreatic cancer (Supplementary Table 13). Additionally, the adjustment decreased the statistical significance of Shannon diversity (Figure 4D), which is one of the most common characteristics to decrease in individuals with diseases^71,72^, in all of the 11 diseases significantly associated with it before the adjustment. In four diseases (ulcerative colitis, ankylosing spondylitis, IBS-D, and slow transit constipation), associations with Shannon diversity were not significant after the adjustment (FDR > 0.05). Overall, our results suggest that microbial loads could confound a substantial portion of disease-associated gut microbial species.

Finally, when deriving absolute abundances of microbial species by taking into account the predicted load, we found that quantitative species profiles reduced biases in relative abundance profiles, therefore reducing over- or underestimation of the significance of species in several diseases associated with microbial load (Supplementary Figure 14).

## Discussion

In this study, we developed novel machine-learning models to predict microbial loads solely based on the relative species and gene abundances of the fecal sample (Figure 1). The benchmarking (cross-validation within the training cohorts and application to independent study populations) as well as the consistency with existing knowledge on microbial load changes (e.g. after antibiotic treatment) supported the robustness of the prediction. Although various methods are available to experimentally quantify microbial load in fecal samples (e.g. flow cytometry, qPCR, and spike-in DNA), the present models are a convenient way to predict load without additional wet lab assays, particularly for existing public fecal metagenomes, as it can be directly inferred from relative microbiome profiles.

Application of the prediction models to the large-scale microbiome datasets uncovered various host and environmental factors significantly associated with predicted microbial loads, including age, sex, diet, diseases, and medications (Figure 2). Although many of these factors are interdependent, microbial load appears as a major factor that could explain indirect associations whose mechanisms were unknown (e.g. higher microbial diversity for elderly people and females). Our analysis also revealed significant differences in microbial loads across various diseases (Figure 3), which indicates that patient microbiomes are not only affected directly by the disease, but also indirectly by physiological (e.g. water content, oxygen concentration, and pH) and physical (e.g. transit time) changes that accompany the disease. For example, diarrhea is common in various gastrointestinal and infectious diseases^73–75^, while constipation is a common complication for several neurological diseases such as Parkinson’s disease^76^, Alzheimer’s disease^76^, and multiple sclerosis^77^, and a risk factor for colorectal cancer^78^. Furthermore, drug treatment could change bowel movement and induce constipation (e.g. opioids, antipsychotics, and non-steroidal anti-inflammatory drugs [NSAIDs])^79^. These observations and findings that more than half of the disease-microbe associations lost their significance after the adjustment in several diseases (Figure 4), suggest that the microbial load can be a major confounder in disease association studies. This may partly explain the reason why the microbial signatures of a particular disease are often non-specific and shared across multiple diseases, as observed in previous meta-analysis studies^65,80,81^. Although various interconnected factors shape microbial load (e.g. transit time, diarrhea/constipation, and inflammation), determine the gut microbial composition in general, and confound disease signatures in particular, the predicted microbial load is useful in disentangling such confounding effects in microbiome studies. By incorporating the fecal microbial load as a baseline factor and excluding associated species, we might better identify gut microbes associated with a disease and develop biomarkers with improved specificity.

The machine-learning model that predicts fecal microbial loads, exclusively based on the relative species abundances, is freely available (MLP: Microbial Load Predictor, https://microbiome-tools.embl.de/mlp/). Although the model was accurate enough to capture known and unknown biological associations, its accuracy will likely increase through refinement with more data or better machine-learning algorithms. In principle, the approach can also be applied to other habitats, making microbial loads comparable, for example enabling important global studies, such as better estimates of biomass on Earth.

## Acknowledgments

We thank members of the Bork group at EMBL for their support and constructive discussions. We also thank Anna Głazek, Anna Schwarz, Roman Thielemann, Leonie Thomas, Ela Cetin, Moritz von Stetten, Mariam Hassen, and Kasimir Noack for their help with the metadata curation. This work was supported by funding from the European Union’s Horizon 2020 research and innovation program under grant agreement numbers 668031 (GALAXY) and 825694 (MICROB-PREDICT). This reflects only the author’s view, and the European Commission is not responsible for any use that may be made of the information it contains. The study was also supported by the Novo Nordisk Foundation through a Challenge Grant “MicrobLiver” (grant number NNF15OC0016692) and through core grant (grant number NNF18CC0034900), the Innovation Fund Denmark (grant number: 0603-00484B), by the EMBO Installation Grant (No. 3573), Estonian Research Council Grant (PRG1414), and the Deutsche Forschungsgemeinschaft (DFG, German Research Foundation) – project number 460129525 and 403224013 (project A09). S.N. was partially supported by the Overseas Postdoctoral Fellowships of the Uehara Memorial Foundation. C.E.F. was supported by the BRIDGE – Translational Excellence Programme (grant number: NNF18SA0034956), Steno Diabetes Center Sjaelland, and The Region Zealand Health Scientific Research Foundation. N.N was partially supported by the Japan Agency for Medical Research and Development (AMED) (Research Program on HIV/AIDS: JP22fk0410051, and Research Program on Emerging and Re-emerging Infectious Diseases: JP22fk0108538), and the Ministry of Health, Labour, and Welfare, Japan (grant number:22HB1003).

## Author contribution

Conceptualization, S.N., M.K., P.B.; methodology, S.N.; data analysis, S.N., O.A., M.I.K., A.F., C.S., M.R.; sample collection and sequencing, J.K.H, L.A.H, M.I., R.S., N.T., M.A., F.B., C.B., C.E.F., J.H., T.N., J.S.P., M.S.T., J.T., N.N., E.O., A.K.; cell counting, E.S., H.B.J., T.H.; collection of public metagenomes and metadata curation, T.S.B.S., P.F., S.N, A.N.F. M.K.; writing, S.N., M.K., P.B.; supervision, J.T., E.O., A.K., T.H., M.K., P.B. All authors discussed the results, reviewed the manuscript, and approved the final manuscript.

## Consortia

**GALAXY**: Manimozhiyan Arumugam, Peer Bork, Torben Hansen, Roland Henrar, Hans Israelsen, Morten Karsdal, Cristina Legido-Quigley, Hans Olav Melberg, Maja Thiele, Jonel Trebicka, and Aleksander Krag (coordinator).

**MicrobLiver**: Peer Bork, Mathias Mann, Jelle Matthijnssens, Aleksander Krag, and Torben Hansen (coordinator).

## Methods

### GALAXY/MicrobLiver study population

A total of 1,906 fecal samples were collected in the GALAXY/MicrobLiver study population. These samples were derived from 9 different cohorts (GALA-ALD^82–84^, GALA-HP^85,86^, GALA-RIF^87,88^, AlcoChallenge^89–92^, HCO^93,94^, GALA-POSTBIO, GastricBypass, The HOLBAEK Study (HOLBAEK)^95^, and TIPS^96–99)^ with different study designs and objectives. These studies involved diverse participant groups, including healthy individuals (GALA-HP), patients with chronic alcohol-related liver disease (ALD) (GALA-ALD and TIPS), those with severe obesity (GastricBypass), those born with low birth weight (HCO), children and adolescents with obesity (HOLBAEK), and patients with dietary (GALA-POSTBIO), alcohol (AlcoChallenge), and drug interventions (GALA-RIF). Out of the 1,906 samples, 12 samples were excluded as outliers from the downstream analyses due to having substantially lower microbial loads than other samples (<10% of the median value of other samples). In total, 1,894 fecal samples from 1,351 participants were used in the study (Supplementary Tables 1 and 2). The objective of each cohort, study design, inclusion, and exclusion criteria were described as follows.

### GALA-ALD

This is a prospective, single-center, biopsy-controlled, cross-sectional study covering the full range of alcohol-related liver disease (ALD)^82–84^. Patients were recruited between 2013 and 2018 in the Region of Southern Denmark. Inclusion criteria comprised individuals aged 18-75 years with prior or current chronic alcohol overuse, which was defined as more than 24 g/day for women and more than 36 g/day for men for over a year, and informed consent to a liver biopsy. Exclusion criteria included solid evidence of cirrhosis, concurrent liver diseases, severe illnesses with less than 12 months expected survival, contraindications to percutaneous liver biopsy, severe alcohol-related hepatitis, hepatic congestion or bile duct dilation as shown by ultrasound, HIV positive status, ongoing substance abuse other than alcohol, and inability to comply with the study protocol. Participants were sourced from both primary and secondary healthcare, encompassing populations at low versus moderate-high prevalence of cirrhosis.

### GALA-HP

This longitudinal study involved healthy participants recruited between 2016 and 2018 at Odense University Hospital in Denmark. The inclusion criteria specified individuals aged 18-75 who were matched by sex, age and (partially) BMI to patients from the GALA-ALD study^85,86^. The exclusion criteria included current alcohol consumption exceeding 7 units per week, prior harmful alcohol use, known liver disease, elevated liver enzymes or altered liver function tests, signs of altered glucose metabolism, signs of other metabolic diseases, infection/inflammation, significant vitamin/mineral deficiencies, any known chronic diseases, ongoing substance abuse, use of any medication (besides infrequent use of mild pain relievers), and use of antibiotics within the last six months. Stool samples were collected at home, frozen in the home freezer immediately, and brought to our unit (with cooling elements to remain frozen) for –80 °C storage within 24 hours.

The study protocol for the GALA-ALD and GALA-HP was approved by the ethics committee for the Region of Southern Denmark (nos. S-20160006G, S-20120071, S-20160021 and S-20170087) and is registered with both the Danish Data Protection Agency (nos. 13/8204, 16/3492 and 18/22692) and Odense Patient Data Exploratory Network (under study identification nos. OP_040 and OP_239 [open.rsyd.dk/OpenProjects/da/openProjectList.jsp]). These studies were conducted according to the principles of the Declaration of Helsinki, and oral and written informed consent was obtained from all participants.

### GALA-RIF

The GALA-RIF trial, an investigator-initiated, randomized, double-blind, placebo-controlled, single-center, phase 2 study, was conducted to evaluate the efficacy of rifaximin-α in patients diagnosed with alcohol-related liver disease through liver biopsy^87,88^. Patients were allocated in a 1:1 ratio to either rifaximin-α or placebo for 18 months. Patients were recruited at the Department of Gastroenterology and Hepatology at Odense University Hospital in Denmark. Ethical approval was granted by the regional ethics committee (S-20140078), and the study adhered to the International Conference on Harmonization Good Clinical Practice guidelines, with external monitoring by the Good Clinical Practice Unit at Odense University Hospital. Participants were identified from a cross-sectional study (GALA-ALD) focusing on alcohol-related liver disease. Alcohol overuse was defined as a daily intake of 24g or more for women and 36g or more for men for at least a year. The study excluded patients with a history of hepatic decompensation or any known liver disease. Following the screening, patients at risk of liver fibrosis underwent liver biopsy. From these, patients aged 18–75 years with liver fibrosis and histological features of alcohol-related liver disease were included in the study. Stool samples analyzed in this study were derived from baseline, 1 month, and 18 months (at the end of treatment). EudraCT, number: 2014–001856-51.

### AlcoChallenge

This clinical study aimed to investigate the acute impact of alcohol consumption on the intestine with the hypothesis that acute alcohol intake increased intestinal permeability and inflow of bacterial products to the liver^89–92^. Participants aged 18-75 who met the criteria for ALD, metabolic dysfunction-associated steatotic liver disease (MASLD), or healthy controls were included. Patients with other known causes of liver disease, total alcohol abstinence or desire for it, insulin-dependent diabetes mellitus, cirrhosis, pregnancy, recent antibiotic treatment, liver cancer, severe comorbidities, or inability to follow instructions were excluded. Participants were asked to maintain their habitual diet and alcohol consumption until two days before the alcohol intervention. On the investigation day, participants were fasting and abstained from alcohol for 48 hours. Stool samples were collected by the participants within 24 hours of each visit. Participants were given instructions and material for sample collection. The samples were collected in sealed test tubes and stored immediately in the participants’ freezer. The samples were transported to the hospital as cold as possible using a cooler bag and cooling elements. Upon arrival at the hospital, the samples were stored in a –80 °C freezer. The study was approved by the Ethical Committee of Southern Denmark (S-20160083) and registered at ClinicalTrials.gov (NCT03018990).

### HCO

This study aims to investigate whether 12 weeks of exercise training can revert and/or minimize the deleterious cardiometabolic effects of 4 weeks of carbohydrate overfeeding in individuals born with low birth weight and increased risk of developing type 2 diabetes. This study recruited healthy Caucasian males, born between 1979 and 1980 at full term (gestational weeks 39–41)^93,94^. Exclusion criteria for participants included having diabetes in their first-degree relatives, any chronic or acute diseases, medication intake that could affect the study’s outcomes, BMI > 30 kg/m^2^, physical activity >10 hours per week, alcohol consumption exceeding the national recommendations, and significant weight changes (>2 kg) in the past 6 months. Feces were sampled at home and immediately stored in the freezer at –18 °C or cooler. The samples were picked up by the staff and transported on dry ice to the laboratory and stored at –80 °C. The study was conducted in compliance with the Declaration of Helsinki II and approved by the ethical committee of the Capital Region of Denmark, with identifier H-4-2014-128. The research has been registered under the ClinicalTrials.gov identifier: NCT02982408. All participants provided written informed consent to participate in the study.

### GALA-POSTBIO

A 24-week prospective, randomized controlled clinical trial aiming to investigate if a postbiotic drink made of fermented oats, ReFerm®, could alter the progression of liver disease compared to an active comparator, Fresubin®. From March 2019 to January 2021, 56 patients were recruited and included in the study. The trial was held at the Department of Gastroenterology and Hepatology at Odense University Hospital in Denmark. Ethical approval was granted by the regional ethics committee (S-20170163) and the Danish Data Protection Agency (19/6646). Patients were allocated in a 1:1 ratio to either ReFerm® or Fresubin® treatment groups. Clinical investigations were conducted at baseline, 4 weeks, 24 weeks (end of intervention), and after a wash-out period of 6 to 8 weeks. Inclusion criteria were outpatients with stable, compensated advanced chronic alcohol-related liver disease between 30 and 75 years. Compensated advanced chronic alcohol-related liver disease was defined as liver stiffness ≥15 kPa or a newly performed (<6mdr) liver biopsy with Kleiner Fibrosis Stage ≥ 3 or a liver biopsy 6 months with Kleiner Fibrosis Stage ≥ 3 and a current liver stiffness ≥10 kPa. Eligible patients had a prior or ongoing harmful alcohol intake defined as an average of ≥24g alcohol/day for women and ≥36 g/d for men for ≥ 5 years. Exclusion criteria were Child-Pugh C score, Meld-Na ≥15, hospitalization within three months, moderate or severe ascites, high-risk varices needing interventional treatment, known liver disease other than alcohol-related, antibiotic treatment in the prior three months, and treatment with nutritional drinks, probiotics or prebiotics within the last three months. ClinicalTrials.gov ID: NCT03863730.

### GastricBypass

The bariatric study cohort is based on 70 patients with a BMI > 35.0 kg/m2 undergoing laparoscopic bariatric surgery (either Roux-en-Y gastric bypass (n=30) or sleeve gastrectomy (n=40)). The design is best described as a prospective cohort study. Study subjects were included between December 2016 and September 2019 at Copenhagen University Hospital Hvidovre. The study subjects fulfilled the existing criteria for bariatric surgery issued by the Danish Health Authorities (BMI>35.0 kg and metabolic comorbidity and/or arthrosis in lower extremities OR BMI>50 with or without metabolic comorbidity/arthrosis in lower extremities), including a mandatory weight loss of 8% before surgery. The mode of surgery (Roux-en-Y gastric bypass or sleeve gastrectomy) was decided by the endocrinologists at the Endocrinology Department. Study-specific exclusion criteria were current or previous alcohol consumption of 2.5 units/day for men and > 1.5 units/day for women, use of antibiotics within one month prior to surgery, preexisting liver disease other than metabolic dysfunction-associated steatotic liver disease, pre-existing disease in the lipid metabolism and acute or chronic inflammatory disease, or an ethnic origin other than North European. On the day of surgery (aka baseline visit) fasting project blood samples were collected. The fecal samples were collected 1-7 days prior to surgery and immediately frozen. During surgery, liver and adipose tissue were sampled. Follow-up visits including collection of fecal-and blood samples were conducted three, six, and 12 months after surgery. Fecal samples at the baseline, three, and 12 months were analyzed in this study. The study protocol was approved by the Regional Scientific Ethics Committee (H-16030784 and H-16030782). Written and oral informed consent was obtained from all study participants. The study was conducted according to the Declaration of Helsinki.

### The HOLBAEK Study

We collected fecal samples from 397 5–19-year-olds of which 331 were from a hospital-based obesity clinic cohort and 66 were from a population-based cohort. The hospital-based obesity clinic cohort consists of children and adolescents enrolled in multifaceted obesity management from January 2008 onwards at a hospital-based obesity clinic^95^. These patients were referred from general practitioners, pediatric departments, or community-based doctors. In the hospital-based obesity clinic cohort, the longitudinal data collection began just prior to the initiation of non-pharmacological obesity treatment and continued with the subsequent contacts in the clinic in a systematic, family-based, person-centred, chronic care setting. The only inclusion criterion was a referral to the hospital-based obesity clinic. Importantly, no a priori age- or other exclusion criteria would make a child or adolescent ineligible for treatment or inclusion in the clinic. The population-based cohort consists of children and adolescents recruited from October 2010 onwards without selection pertaining to body weight or BMI. Recruitment took place at schools and high schools across 11 municipalities in Region Zealand and the Capital Region in Denmark. All children and adolescents at the participating schools were considered eligible for inclusion regardless of age, and no exclusion criteria were applied. Informative recruitment meetings for potential participants were held during school hours and written material was delivered to the parents. Stool samples were collected at participants’ homes, immediately frozen in their home freezers, and then transported to the laboratory with cooling elements to ensure they remained frozen. Upon arrival, the samples were stored in freezers at a temperature of –80 °C within 24 hours of collection. The HOLBAEK Study was approved by the Ethical Committee of Region Zealand (Project number: SJ-104), The Danish Data Protection Agency (REG-043-2013), and other collateral project approvals and was registered at ClinicalTrials.gov on June 26, 2009 (NCT00928473). All procedures in relation to the biobank are performed in accordance with the Helsinki Declaration. Written informed consent was obtained from parents/legal guardians or from the adolescents themselves when above the age of 18 years.

### TIPS

The TIPS study is a single-center prospective study in patients with decompensated cirrhosis who received a transjugular portosystemic shunt as part of the NEPTUN study (NCT03628807) at the Department of Internal Medicine I, University Clinic Bonn (Germany)^96–99^. For this study, stool samples from 84 patients were obtained between 2014 and 2018. The mean age was 58 years (range 18-84 years), 53% of the patients were male and the majority of patients had alcohol-induced cirrhosis (n=62), followed by viral hepatitis (n=8) and other etiologies (n=18). The stool samples were collected during the inpatient treatment of the patients shortly before the TIPS procedure and stored directly at -80 °C degrees until further use. The study was approved by the local ethics committee of the University of Bonn (029/13), and all patients signed an informed written consent in accordance with the Helsinki Declaration.

### DNA sequencing of fecal samples

Microbial DNA was extracted from collected stool samples using Qiagen AllPrep PowerFecal DNA/RNA Kit (Qiagen, Hilden, Germany) following the manufacturer’s protocol in the GALA-RIF, AlcoChallenge, HCO, GALA-POSTBIO, and TIPS cohorts. The same protocol, except for an additional phenol-chloroform extraction step after the step of lysing microbial cells, was used in the GALA-ALD, GALA-HP, HOLBAEK, and GastricBypass cohorts. Metagenomic sequencing libraries were prepared using the NEBNext Ultra II DNA Library Prep kit (New England Biolabs, MA, USA) with a targeted insert size of 350-400bp and Dual Index multiplex oligos. Libraries were prepared using a liquid automated system (Beckman Coulter i7 Series) and sequenced on an Illumina HiSeq 4000 platform (Illumina, San Diego, CA, USA) with 2x150bp paired-end reads.

### Quality control of sequenced reads

Sequenced reads were processed to remove low-quality reads and host-derived reads using ngless (v1.1)^100^. Nucleotide calls with a Phred quality score of less than 25 were removed from the 3’ end and reads less than 45 nucleotides long after the removal were discarded. Reads representing human DNA were identified by comparing all reads’ sequence similarity to the human reference genome (hg38). Any reads with greater than 90% similarity to the human genome were discarded. After this quality control, reads were re-classified as paired or as singles, where, respectively, both or only the forward and reverse reads are present in the final dataset.

### Cell counting

Bacterial cell counting was performed as previously described^101^. Briefly, frozen (-80°C) fecal samples were diluted, mechanically homogenized and afterward fixed with 2% Paraformaldehyde (10 min, RT; VWR). To minimize clumps, the samples were filtered through a cell strainer. The resulting bacterial cell suspension was stained with SYBR Green I (1:200,000 (Fisher Scientific), in DMSO (Sigma-Aldrich)) and incubated in the dark for 30 min. Measurements were performed at a pre-set flow rate of 0.5 μL/sec, and a known concentration of 123count eBeads (Invitrogen) was added for accurate bacterial cell count estimation. Measurements were performed using a BD Fortessa LSRII flow cytometer (BD Biosciences) (GALA-HP, Alcochallenge, GALA-ALD, TIPS, HCO cohorts) and BD Fortessa 3 flow cytometer (BD Biosciences) (GALA-POSTBIO, GALA-RIF, HOLBAEK, GastricBypass cohorts), and data were acquired using BD FACSDiVa software. A collection threshold value of 200 was applied on the FITC (530/30 nm) channel. Fluorescence intensity was collected at green (530/30 nm, FITC), blue (450/50 nm, Pacific Blue), yellow (575/26 nm, PE), and red (695/40 nm, PerCP-Cy5.5) fluorescence channels. Forward- and side-scattered (FSC and SSC) light intensities were also collected. Data was processed in R using the flowcore package (v1.11.20)^102^ in R Studio (v4.1.2). Fixed gating strategy was applied for all samples to allow direct comparison between measured samples. Bacterial cell counts, estimated from pre-set flow rate, were adjusted with internal control counts, included on each plate, to correct for batch effects.

### Taxonomic and functional profiling of metagenomes

Species-level taxonomic profiles of the samples were obtained with the marker-gene-bases method using mOTUs (v2.5)^36^. Functional profiles were obtained by mapping metagenomic reads to the sub-catalog of the human gut microbiome in the global microbial gene catalog (GMGC)^37^ using BWA-MEM (v0.7.17)^103^ with the default parameters. The genes were functionally annotated using eggNOG-mapper (v1.0.3)^104^ against eggNOG database 5.0^105^ and KEGG orthologies^106^ were assigned to each gene. The number of reads mapped to each KEGG orthology was counted using gffquant (v2.9.1) (https://github.com/cschu/gff_quantifier) where the count of the number of reads aligning multiple genes was distributed to each gene by dividing by the number of the genes.

### MetaCardis dataset

Fecal metagenomes from the MetaCardis project (n = 1,820)^33–35^ were downloaded from the European Nucleotide Archive under the accession numbers PRJEB41311, PRJEB38742 and PRJEB37249. Microbial load data for these samples were obtained in the study of Forslund *et al*^34^. Out of 1820 samples, eight samples were excluded from the downstream analyses as outliers due to significantly lower microbial loads than other samples (9.7E+09 and 1.1E+11, respectively). In total, 1,812 samples were used in the following analyses. Taxonomic and functional profiles of the microbiomes were obtained with the same procedure described above.

### Association analysis between the gut microbiome and microbial loads

To investigate correlations between the microbiome profile (i.e. species-level taxonomic and functional compositions) and the experimentally measured microbial load, Pearson correlation coefficients were calculated between the log10 transformed relative abundance of each microbial species/functions and the microbial load in each cohort separately. Additionally, the analysis was also performed for diversity indexes of the taxonomic profiles such as Shannon diversity, species richness (i.e. the number of detected species), and Simpson diversity. The over-representation of KEGG pathways in the positively- and negatively-correlated functions was identified with the gene set enrichment analysis using the GSEA function in the clusterProfiler package (v4.8.3)^107^. Multidimensional Scaling (MDS) analysis was performed using the metaMDS function in the vegan package (v2.6.4), based on a Euclidean distance matrix derived from log10-transformed relative abundance data with half of the minimum non-zero value as pseudocounts. Enterotypes (i.e. *Bacteroides*, *Prevotella,* and Firmicutes types)^38,39^ of the gut microbiome were determined as described previously (Keller MI *et al*, in preparation) using the pam3 model and they were plotted into the MDS ordination using the envfit functions in the vegan package.

### Construction of prediction models

To construct machine-learning models to predict the microbial load, we employed the eXtreme Gradient Boosting (XGBoost) algorithm^46^, available in the xgboost R package (v1.7.5.1). Prior to model training, we performed unsupervised feature filtering on the species-level taxonomic profiles of the microbiome to exclude minor species (those with < 0.1% average abundance or < 10% prevalence). The relative abundances of each species and the microbial loads were then log10 transformed before the training. For the species, we added half of the non-zero minimum values in the dataset to each abundance to avoid log10 transformation of 0 values, and further standardized (i.e. z-score). The models were trained using the train function in the caret R package (v6.0.94)^108^ in the GALAXY/MicrobLiver and MetaCardis datasets separately, employing a 5-times repeated 10-fold cross-validation procedure to maximize the root-mean-square error (RMSE) in the model. The hyperparameters were determined through a grid search. For internal validation, we calculated the average predicted microbial loads across the test datasets for each sample and compared these with the actual microbial loads. For external validation, we applied the GALAXY/MicrobLiver model to the MetaCardis dataset, and vice versa, comparing the predicted and actual microbial loads.

### Analysis of 16S rRNA gene data from previous studies

Additional paired data of 16S rRNA gene sequencing and fecal microbial loads were collected from two previous studies^19,24^. For Vandeputte *et al.* 2021 study, the genus-level taxonomic profiles were obtained from the paper. For Vandeputte *et al.* 2017 study, 16S rRNA gene sequencing data were downloaded from the European Nucleotide Archive, under the accession numbers PRJEB21504 and ERP023761. The 16S rRNA gene sequencing data was processed using the DADA2 pipeline and the taxonomic annotation was performed using the RDP training data rdp_train_set_16. A prediction model was constructed based on the data of Vandeputte *et al.* 2021 (n = 707) using the same procedure described above. Then, the model was applied to the other dataset of Vandeputte *et al.* 2017 (n = 95) for external validation.

### Collection of external microbiome datasets

Global dataset: Publicly available human gut metagenomes were downloaded from the European Nucleotide Archive. The dataset was part of a previous study^109^ and was composed of 27,832 samples across 45 countries from 159 studies (Supplementary Table 7). After the downloading, quality filtering was performed using ngless, and bases with <25 Phred quality score were trimmed from the 3’ end, and reads less than 45 bp were excluded. Host metadata such as age, sex, country, antibiotic treatment, and disease were collected from respective study papers manually. Samples from infants, children under 3 years old, and patients who received fecal microbiota transplantation were excluded since their gut microbiomes are substantially different from those of adults^110,111^. Also, samples with a low number of sequenced reads (ie. <1 million) were excluded. Countries were classified into four groups according to the World Bank definition (https://www.worldbank.org/en/home/, accessed in February 2024), which defines high-income, upper-middle, lower-middle, and low-income economies based on gross national income per capita.

Japanese 4D dataset: The Japanese 4D (Disease, Drug, Diet, Daily life) microbiome cohort is a prospective, multicenter, hospital-based cohort established in the Tokyo metropolitan area. A total of 4,198 fecal samples were collected from the participants and processed as described previously^47,112^. Various intrinsic and extrinsic factors (n = 244) were collected from the participants through a combination of self-reported questionnaires, face-to-face interviews, and physicians’ electronic medical records. These factors included anthropometric measurements, lifestyles, dietary habits, physical activities, diseases, and medications (Supplementary Table 8). The protocol for the project was approved by the medical ethics committees of the Tokyo Medical University (approval No.: T2019-0119), National Center for Global Health and Medicine (approval No.: 1690), the University of Tokyo (approval No.: 2019185NI), Waseda University (approval No.: 2018-318), and the RIKEN Center for Integrative Medical Sciences (approval No.: H30-7). All participants provided written informed consent before participation in the project.

Estonian Microbiome dataset: The Estonian Microbiome cohort^113^ is a volunteer-based cohort that currently includes genotyped adults (≥ 18 years old) across Estonia. Fecal samples were collected from 2,509 participants in the cohort and sequenced as described previously^48^. All the participants provided informed consent for the data and samples to be used for scientific purposes. This study received approval from the Research Ethics Committee of the University of Tartu (approval No. 266/T10) and from the Estonian Committee on Bioethics and Human Research (Estonian Ministry of Social Affairs; approval No. 1.1-12/17). Host factors such as anthropometric measurements, lifestyle, diet, disease, and medication were collected from self-reported questionnaires and electronic health records (n = 251, Supplementary Table 8).

### Analysis of the external metagenomic datasets

Species-level taxonomic profiles of the global, Japanese 4D, and Estonian Microbiome samples were obtained with the same method described above. The relative abundance of each microbial species was log10-transformed (1e-4 was added as a pseudo count beforehand) and standardized before the prediction. The MetaCardis prediction model was applied to the profiles and microbial loads were predicted for each sample. The MetaCardis model was employed for the analysis since it was trained on samples derived from more individuals (n = 1,812) than the GALAXY/MicrobLiver model (n = 1,351).

To determine the explained variance of the predicted microbial loads (coefficient of determination) by the collected host factors in the Japanese 4D and Estonian Microbiome cohorts, linear regression analysis was performed using the glm function with the log10 transformed microbial load as a response variable and all the host factors as explanatory variables. Furthermore, the analysis was performed for each metadata category (e.g. lifestyle, diet, medication, and disease) separately, and the explained variance by each category was determined. P-values were adjusted for the multiple comparisons with the Benjamini-Hochberg method^114^.

### Association analysis between diseases and the microbial load

To explore associations between diseases and microbial loads, we performed a meta-analysis of case-control comparisons by combining samples in the global and Japanese 4D datasets. In the global dataset, 58 studies including at least 10 cases and 10 controls were picked up and the case and control samples were collected (Supplementary Table 10). In the Japanese 4D dataset, 9 diseases with >10 patients were selected and age, sex, and BMI matched-controls were defined for each disease using the matchit function of the MatchIt R package (v4.5.3)^115^. Disease-to-control ratios were set to 1:4 when there were enough control samples in the Japanese 4D dataset^116^, while set to 1:1 when the number of samples was insufficient (Supplementary Table 10).

The case and control samples collected above were combined for each disease and a total of 26 diseases with >50 cases and >50 controls were analyzed (Supplementary Table 10). Associations between the diseases and the log10 transformed microbial load were assessed using the glm function with the microbial load as a response variable and the disease condition (i.e. case or control) as an explanatory variable including each study as a covariate. P-values were adjusted for the multiple comparisons with the Benjamini-Hochberg method.

### Association analysis between diseases and the microbiome composition

For the 26 diseases selected, case-control comparisons of the microbiome profiles were conducted. For each disease dataset, species with an average relative abundance of >0.1% and prevalence of >10% were included in the analysis. The relative abundance was log10 transformed after adding half of the non-zero minimum value as a pseudo value. For each microbial species, a linear regression model was applied using the glm function. In this model, the species abundance was included as a response variable and disease condition (i.e. case or control) as an explanatory variable, with each study as a covariate. P-values were adjusted for the number of species for each disease with the Benjamini-Hochberg method^114^.

The set of the obtained effect sizes (beta coefficients) of each species was defined as the microbial signature for the disease. A distance matrix among the 26 diseases was then constructed based on the signature, using Spearman’s correlation coefficient as a distance metric ([1 - Spearman’s correlation] / 2). Then, principal coordinate analysis was performed on this distance matrix using the cmdscale function in the vegan package^117^. The differences in the microbial signatures between the positively- and negatively-associated diseases were examined with permutational analysis of variance using the adonis function in the vegan package with 9,999 permutations for p-value calculation. Additionally, the effect sizes of each disease were compared between the positively- and negatively-associated diseases through a linear regression model, and species discriminating between these two groups were investigated.

To adjust the effect of the microbial load in the case-control comparisons, linear regression models were constructed for each species again by adding the microbial load as a covariate. Effect sizes of each disease on the species and associated P-values were compared between the models with and without the adjustment of the microbial load.

### Comparison of quantitative and relative microbiome profiles in disease association analysis

To explore the advantages of the quantitative microbiome analysis in disease association analysis, we transformed the relative microbiome profiles (RMP) into quantitative microbiome profiles (QMP) (i.e. a profile where species abundances were represented by absolute abundances) by multiplying the relative abundance of each microbial species by the predicted microbial load of the sample. To evaluate the association between each species and disease, the same statistical analyses used for the RMP were performed on the QMP of the same 26 disease datasets. The effect sizes and statistical significance (i.e. p-value) obtained from the analyses were compared with those obtained based on the RMP.

### Data availability

Shotgun metagenomic data sequenced in the GALAXY/MicrobLiver consortia (n = 1,894) are publicly available (upon publication) in the European Nucleotide Archive under the accession number of PRJEB65485 (GALA-ALD), PRJEB65753 (GALA-HP), PRJEB65967 (AlcoChallenge), PRJEB66277 (GALA-RIF), PRJEB67442 (GastricBypass), PRJEB66128 (HCO), PRJEB67944 (GALA-POSTBIO), PRJEB71382 (HOLBAEK), and PRJEB67347 (TIPS). The microbial load of these fecal samples are available in the Supplementary Table 2. Metagenomic data of the Estonian Microbiome Cohort is available in the European Genome-Phenome Archive database (EGAS00001008448). The prediction models we constructed (MLP: Microbial Load Predictor) are available at https://microbiome-tools.embl.de/mlp/ or as R package at https://github.com/grp-bork/microbial_load_predictor/. R codes used to train and construct the prediction models and to generate figures are available at https://github.com/grp-bork/CellCount_Nishijima_2024/.

