## Supplementary Figures for "Fecal microbial load is a major determinant of gut microbiome variation and a confounder for disease associations"

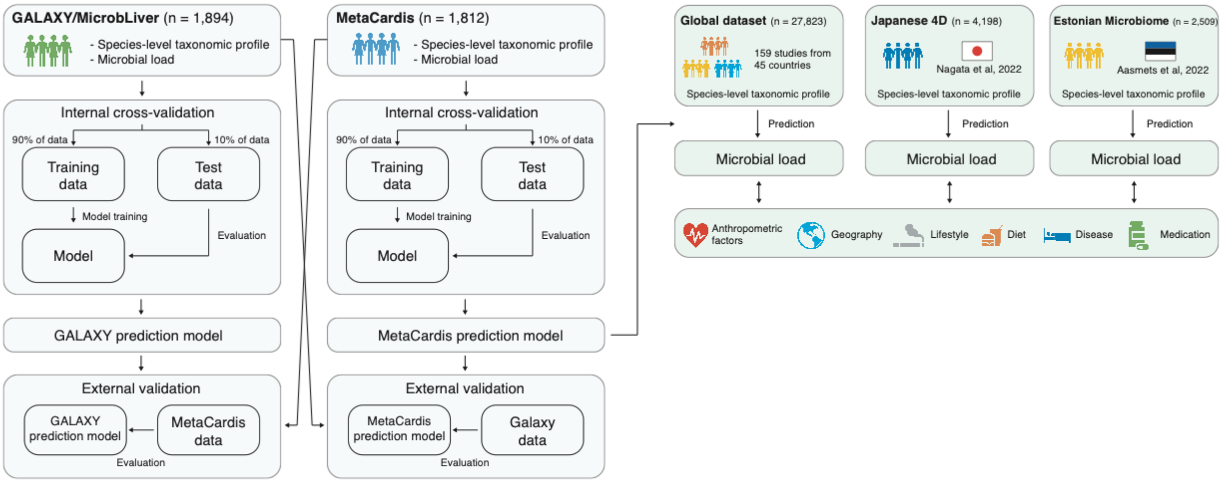


**Supplementary Figure 1 | Flow chart of the analytical processes in this study.**

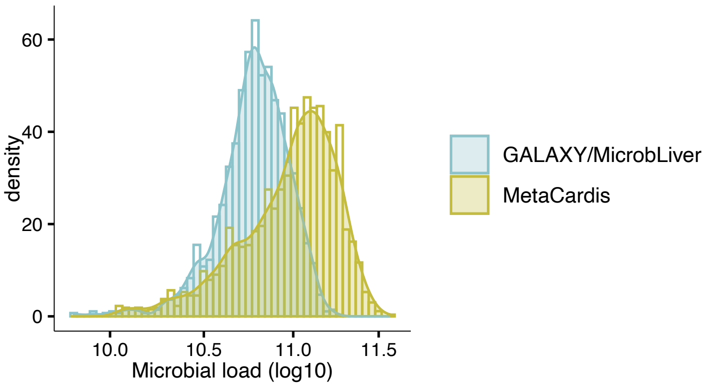


**Supplementary Figure 2 | Comparison of microbial loads between the GALAXY/MicrobLiver and MetaCardis study populations.**
Blue and yellow histograms show experimentally determined microbial loads (total microbial cells per gram) in the GALAXY/MicrobLiver (n = 1,894) and MetaCardis (n = 1,812) study populations, respectively.

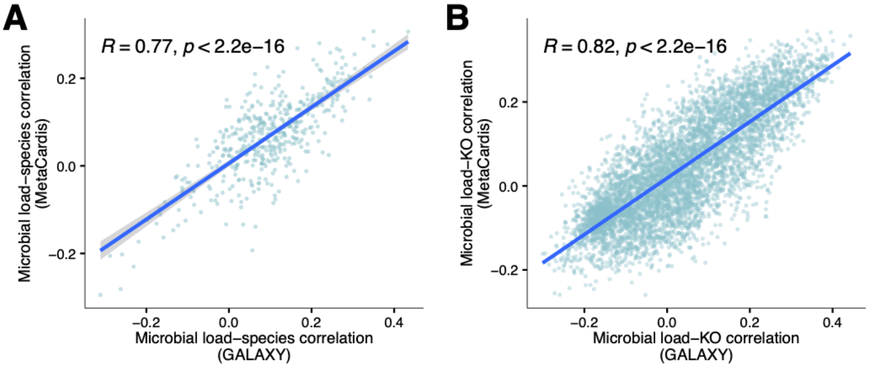


**Supplementary Figure 3 | Comparison of microbiome-load associations between the GALAXY/MicrobLiver and MetaCardis study populations.**

Each circle represents microbial species (**A**) and gene function based on KEGG orthology (**B**). Pearson correlations between log10-transformed relative abundances of species/gene and microbial loads were calculated within the GALAXY/MicrobLiver (n = 1,894) and MetaCardis (n = 1,812) study populations. Then, the correlation coefficients were compared between the two study populations. For the species analysis, those with a mean relative abundance of >0.01% and prevalence of >10% in both studies were included. For the gene function analysis, KEGG orthologies with mean relative abundance of 0.0001% and prevalence of >10% in both studies were included.

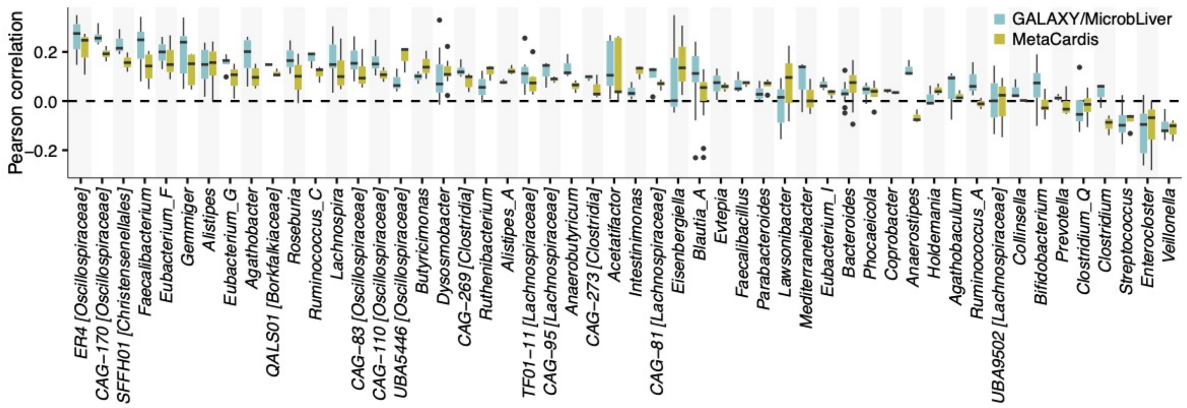


**Supplementary Figure 4 | Species-microbial load correlations summarized at the genus level.**
Pearson correlations were calculated between log10-transformed relative abundances of microbial species and microbial loads in the GALAXY/MicrobLiver (n = 1,894) and MetaCardis (n = 1,812) study populations. The correlation coefficients were summarized at the genus level retrieved from the genome taxonomy database (GTDB). Species with a mean relative abundance of >0.01% and prevalence of >10% in both studies were included in the analysis. The 52 genera containing at least 5 species are shown in the plot.

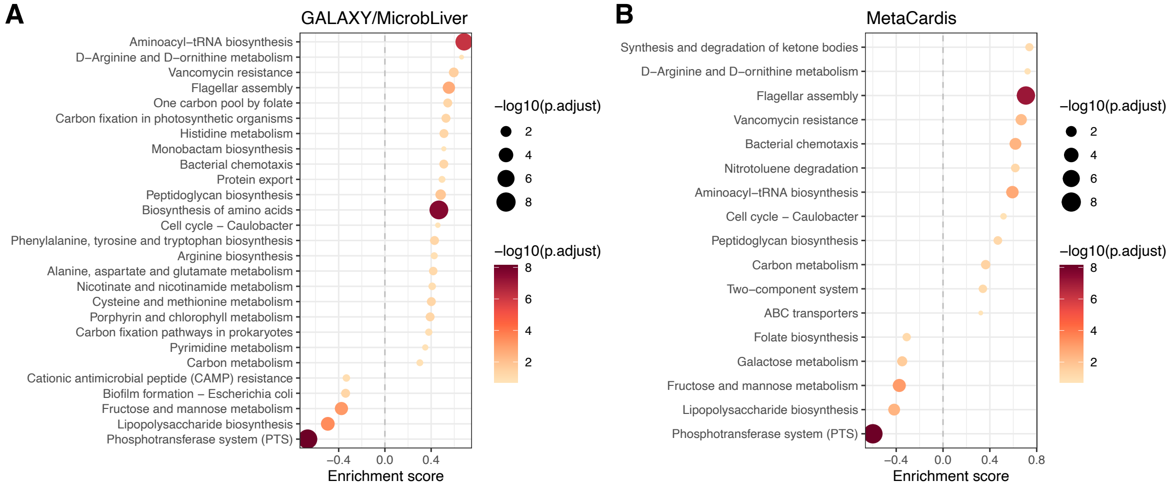


**Supplementary Figure 5 | Pathway enrichment analysis for genes associated with microbial load.** Pathway enrichment analysis was performed to identify over-represented KEGG pathways in positively- and negatively correlated genes with the microbial loads. Pearson correlations were calculated between relative abundances of KEGG orthologies and microbial loads (both log10 transformed) in the GALAXY/MicrobLiver (n = 1,894, **A**) and MetaCardis (n = 1,812, **B**) study populations. Over-representations of KEGG pathways in the positively- and negatively associated genes were assessed using the GSEA function in the clusterProfiler package. Pathways showing statistical significance of FDR < 0.1 are shown in the plot. KEGG orthologies with mean relative abundance of >0.0001% and prevalence of >10% in both studies were included in the analysis.

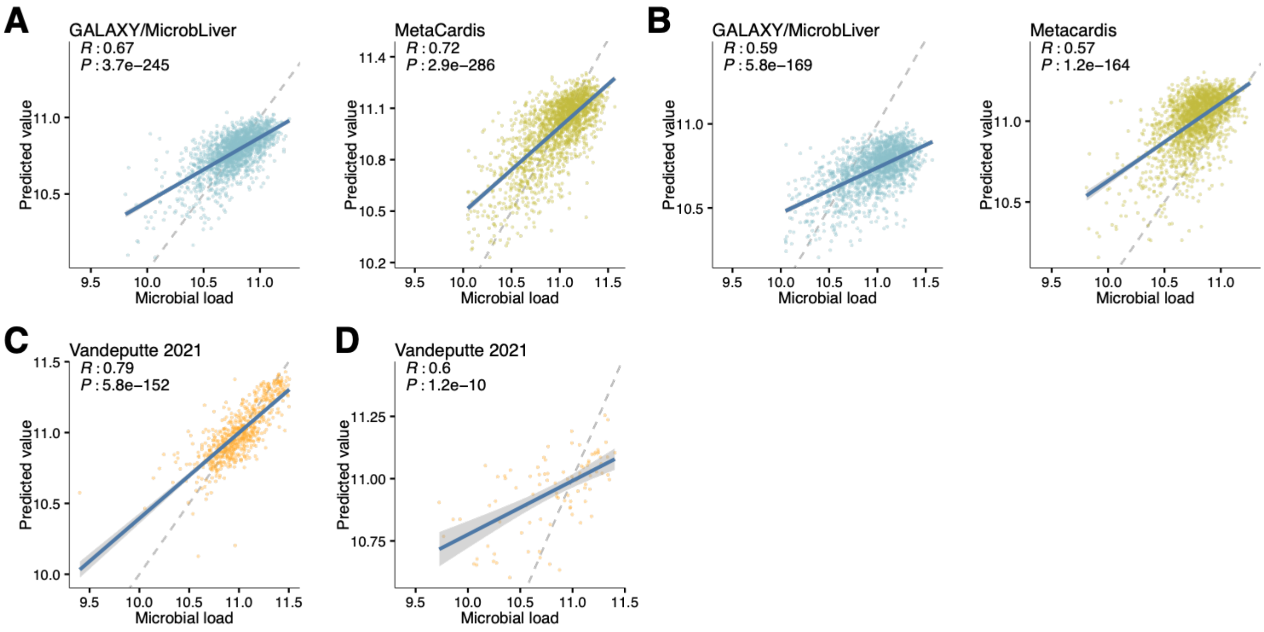


**Supplementary Figure 6 | Internal and external validation of the prediction models based on the functional and 16S rRNA gene profile.**

Evaluation of the prediction models trained on the functional profile of the microbiome (KEGG orthology level) in the GALAXY/MicrobLiver (n = 1,894) and MetaCardis (n = 1,812) study populations (**A, B**) and the 16S rRNA gene profile at the genus level from Vandeputte *et al.* 2021 (n = 707) (**C, D**). The XGBoost algorithm was employed to construct models and they were trained based on relative abundances of KEGG orthology or genera. Internal validation was performed using 5-times repeated 10-fold cross-validation (**A**, **C**). External validation was performed by exchanging the datasets between the GALAXY/MicrobLiver and MetaCardis studies for the functional model (**B**) while employing an external dataset from Vandeputte *et al.* 2017 (n = 95) for the 16S rRNA gene model (**D**).

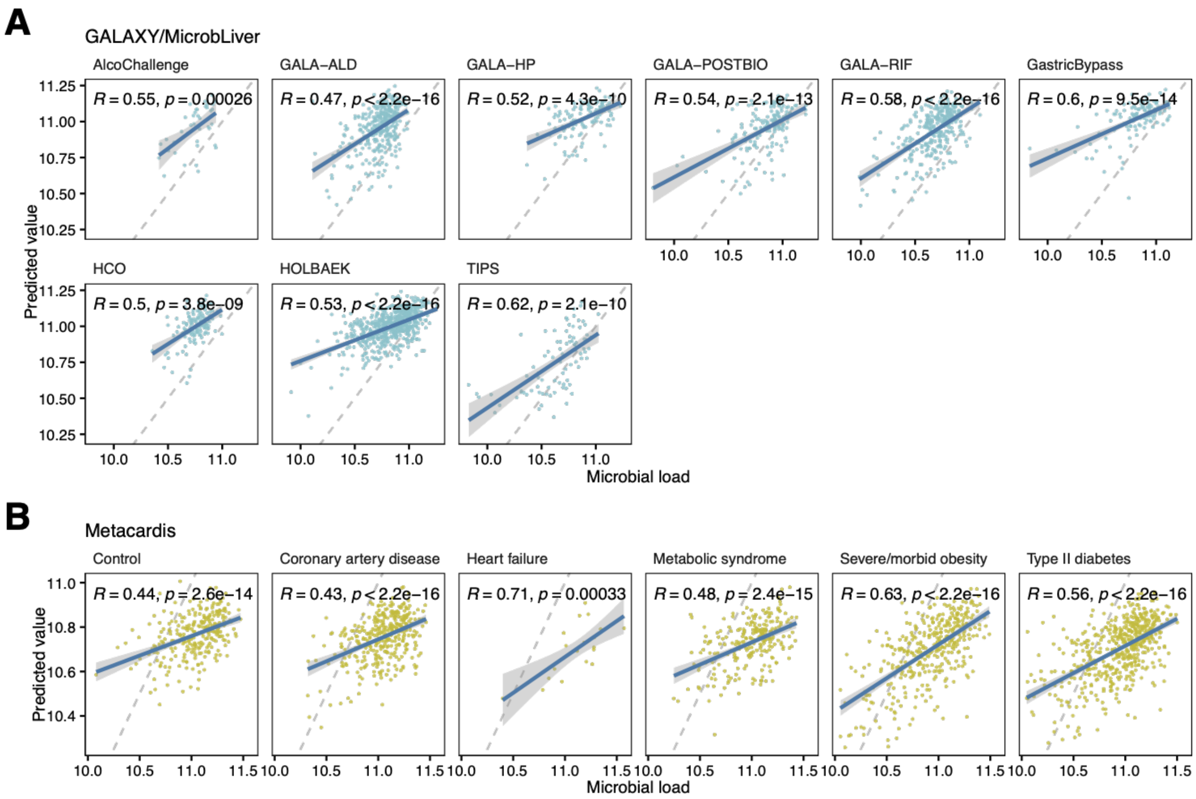


**Supplementary Figure 7 | Robustness of the prediction accuracies across various disease subgroups.**
Consistencies of the prediction accuracies across sub-cohorts in the GALAXY/MicrobLiver study population (**A**) and various disease subgroups in the MetaCardis study population (**B**). Prediction models constructed on the GALAXY/MicrobLiver and MetaCardis were applied to each other's dataset and predicted values were compared to actual microbial loads. Each scatter plot shows prediction accuracy for sub-cohorts in the GALAXY/MicrobLiver study population (AlcoChallenge, n = 39; GALA-ALD, n = 333; GALA-HP, n = 127; GALA-POSTBIO; n = 161, GALA-RIF, n = 317; GastricBypass, n = 129; HCO, n = 125; HOLBAEK, n = 579; TIPS, n = 84) and disease groups in the MetaCardis study population (Control, n = 275; patients with coronary artery disease, n = 361; heart failure, n = 21; metabolic syndrome, n = 246; severe/morbid obesity, n = 373; type II diabetes, n = 536). The solid blue line shows regression lines and the gray dashed lines represent 1:1 reference lines. The shadow of the regression line represents a 95% confidence interval.

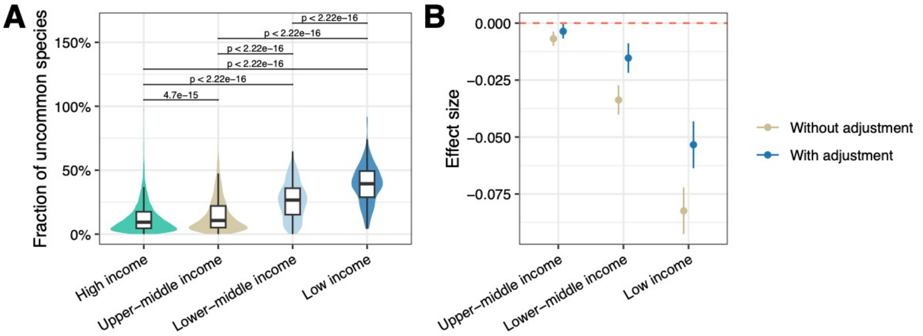


**Supplementary Figure 8 | Evaluation of the model's bias due to the training on only samples from high-income countries.**

Since the prediction model was constructed using samples from high-income countries, we examined whether the difference in the gut microbiomes between high- and low-income countries (e.g. species prevalent in low-income countries but not in high-income countries) confounds differences in the predicted load between them. To assess this, we quantified the total abundance of species not incorporated into the prediction model (i.e. fraction of uncommon species in high-income countries) (**A**) and examined the effect sizes of 4 economic categories on the microbial load with and without adjustment with the fraction using the glm function in R (**B**). **B**, The plots show the effect sizes of each economic category on the microbial load (compared to the high-income country), and the error bars represent 95% confidence intervals. Lower-middle and low-income countries had a significant negative effect on the microbial load, even after the adjustment for the fraction of uncommon species. This result suggests that the difference in the microbiomes between high- and low-income countries is not attributed to differences in microbial load between them.

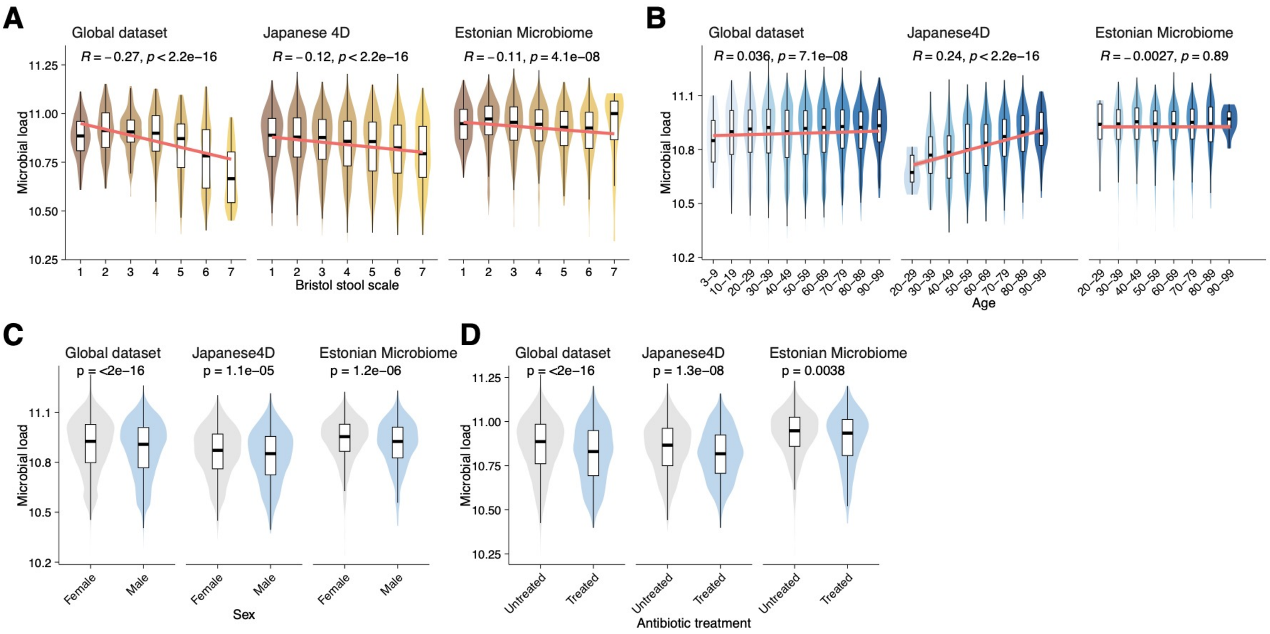


**Supplementary Figure 9 | Predicted microbial loads are significantly associated with various host factors.** Associations between predicted microbial loads and Bristol stool scale (**A**), host age (**B**), host sex (**C**), and antibiotic treatment (**D**) in the global dataset, Japanese 4D, and Estonian Microbiome cohorts. Statistical significance shown in the plots was calculated using Pearson correlation analysis (**A, B**) and Wilcoxon-rank sum test (**C, D**).

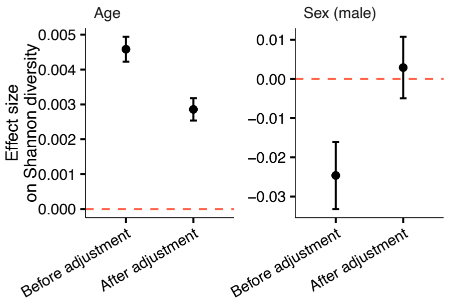


**Supplementary Figure 10 | Microbial loads confound age- and sex-related change in the microbiome.**
Each plot shows the effect size of age (**A**) and sex (**B**) for Shannon diversity of the microbiome in the global and Japanese 4D datasets before and after the adjustment with predicted microbial loads. The effect size was obtained using linear regression analysis including Shannon diversity as the response variable and age and sex as the explanatory variable with and without the predicted microbial loads as covariate. Error bars represent the 95% confidence interval of the effect size.

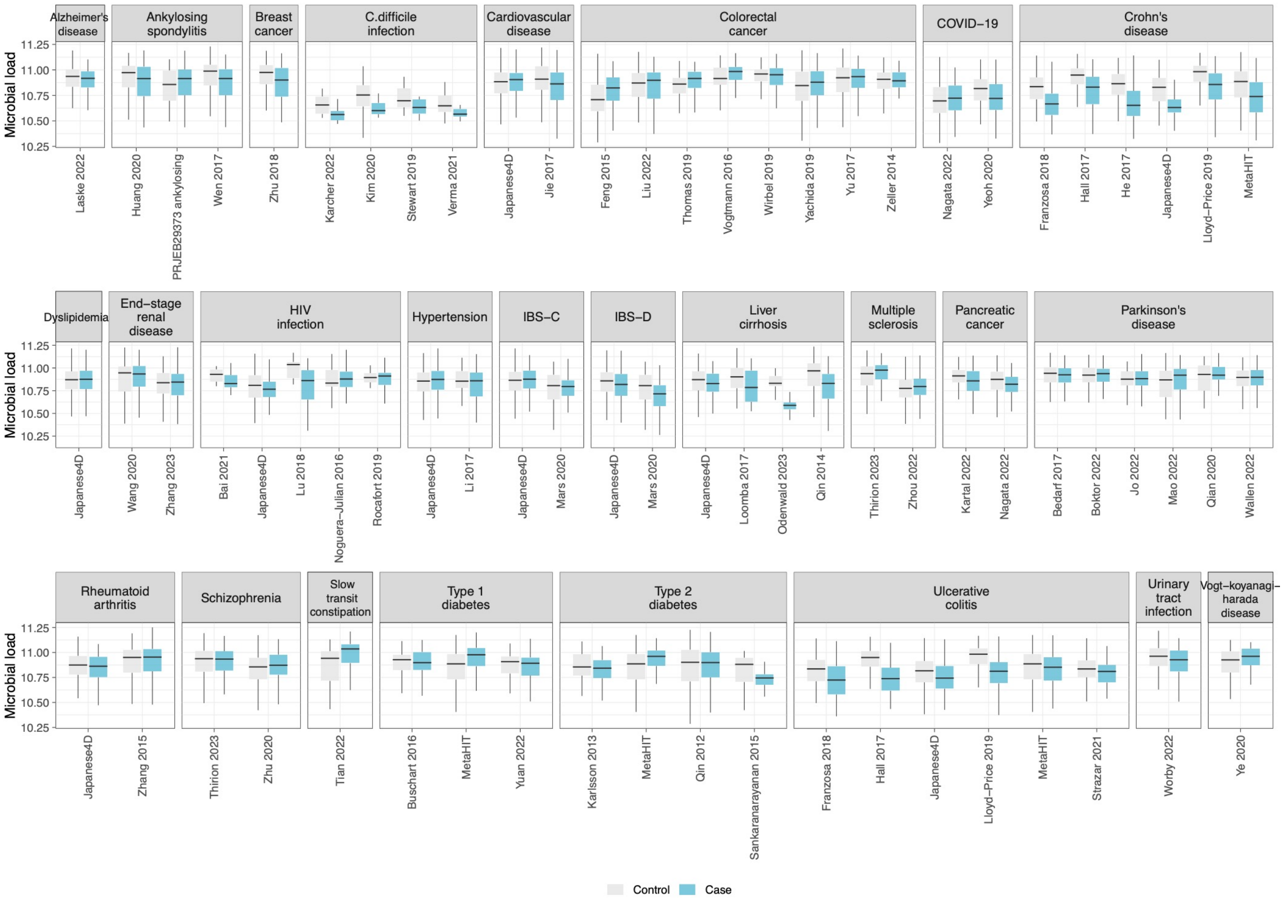


**Supplementary Figure 11 | Comparison of predicted microbial loads between cases and controls.**
Boxplots showing predicted microbial loads (log10 transformed) for cases and controls across 26 diseases. 11,807 cases and 17,118 controls from 58 studies were included in the analysis (Supplementary Table 10).

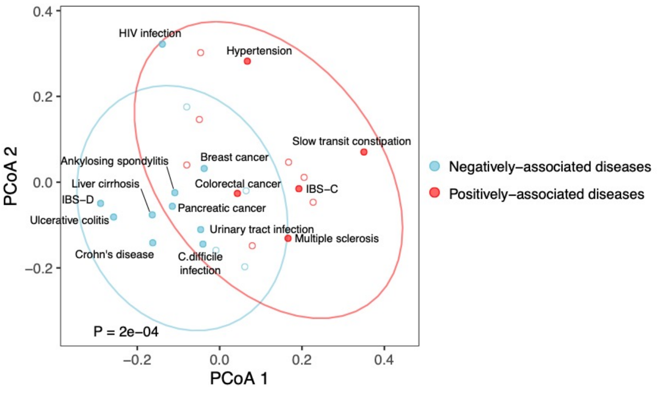


**Supplementary Figure 12 | Comparison of the microbial signature between the positively- and negatively-associated diseases.**

Each circle represents microbial signatures for each disease and the blue and red colors show positively and negatively associated diseases, respectively. To define the microbial signature, relative abundances of microbial species were compared between cases and controls for each disease using a linear regression model incorporating different studies as covariates (Methods). Then, the set of effect sizes of each species was defined as the microbial signature of the disease. The similarity of the microbial signature between diseases was calculated using Pearson correlations and they were transformed into distance. Principal coordinate analysis was performed on the distance matrix of the disease signature using the cmdscale function in the vegan package. The p-value was calculated based on permutational analysis of variance using the adonis function in the vegan package.

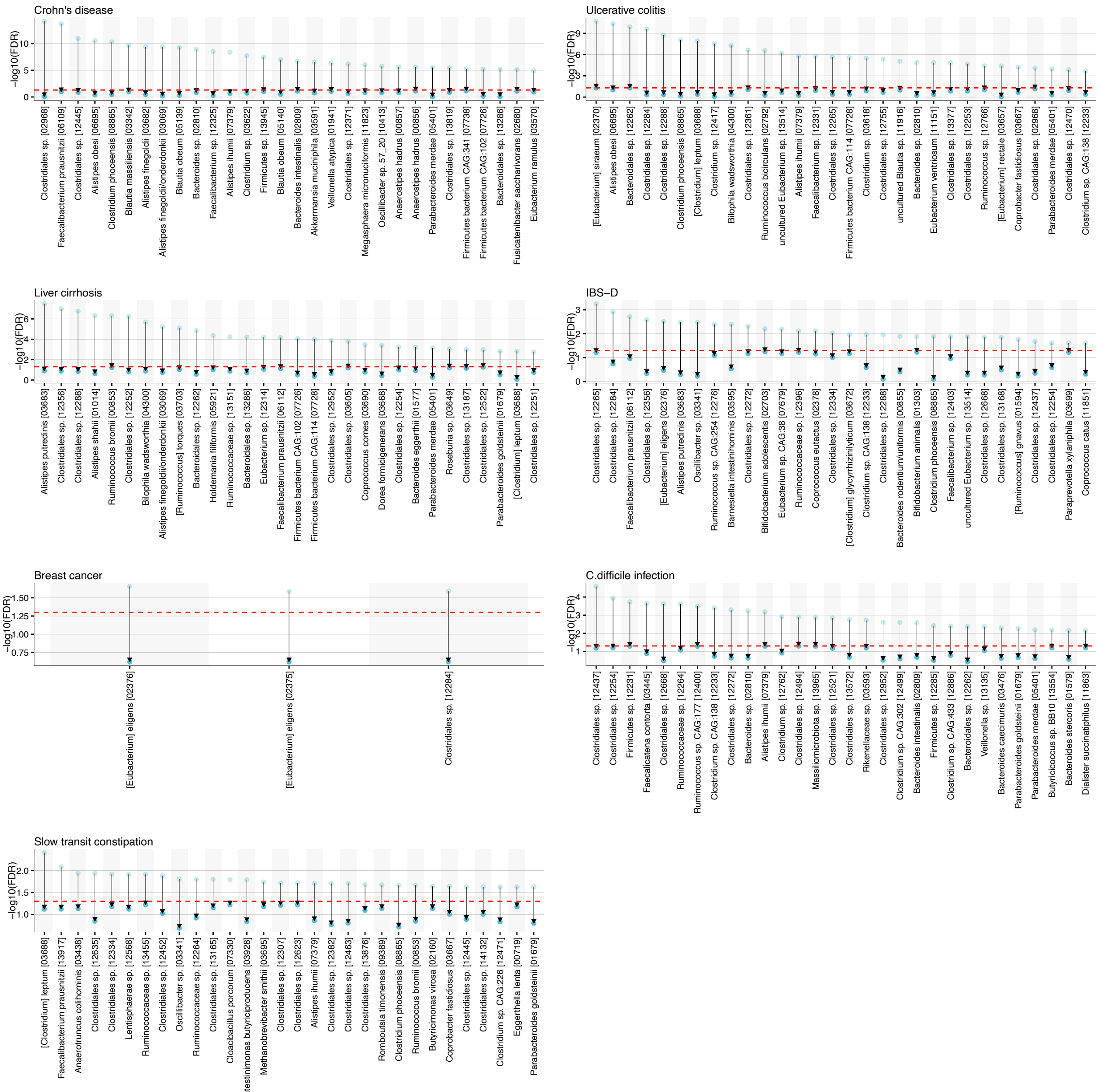


**Supplementary Figure 13 | Microbial load substantially confounds disease-species associations.**
The plot shows species that lost statistically significant associations with diseases after the adjustment with microbial load (FDR > 0.05). Relative abundances of species were compared between cases and controls for each disease and their statistical significance was compared with and without the adjustment. The red horizontal lines show an FDR of 0.05. Arrows represent the changes in the FDR before to after the adjustment with microbial load. Results for the seven diseases, for which the adjustment had the strongest impact, are included in the plot. For diseases with more than 30 species losing significance, the top 30 species with the lowest FDR post-adjustment are shown.

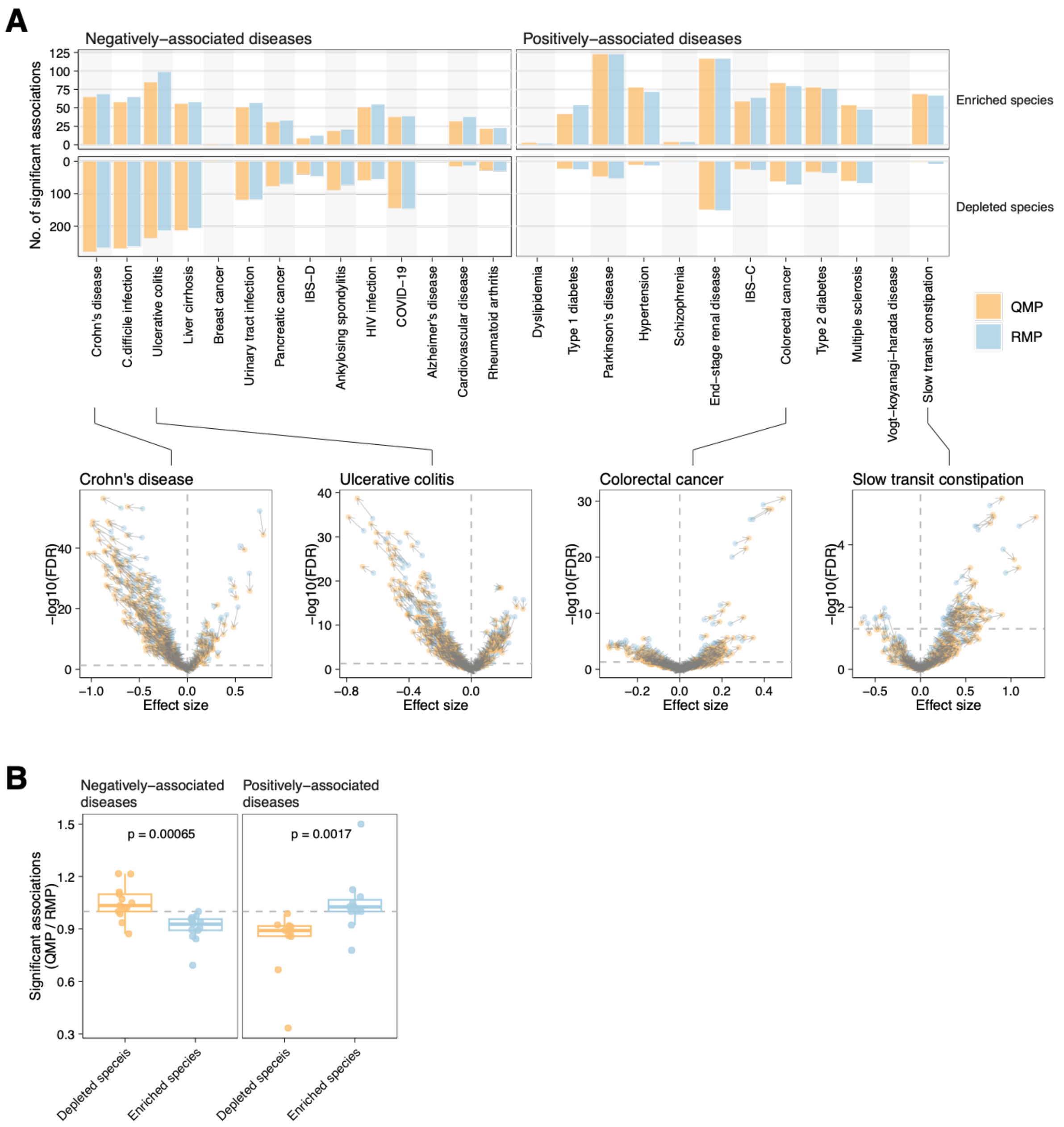


**Supplementary Figure 14 | The predicted microbial loads partially reduce biases derived from the relative nature of the microbiome data**

We compared the results of case-control analyses of each disease between the relative microbiome profile (RMP) (i.e. a profile where species abundances were represented by relative abundance) and the quantitative microbiome profile (QMP) (i.e. a profile where species abundances were represented by absolute abundances by taking into account the predicted microbial load) (Methods). **A**, Bar plot showing the number of significantly enriched and depleted species (FDR < 0.05). The volcano plot below shows the effect sizes and FDR of each species for Crohn's disease, ulcerative colitis, colorectal cancer, and slow transit constipation as examples. Arrows represent the shift of the results from the RMP to QMP. **B**, Boxplot showing the ratio of the number of significant species (FDR < 0.05) between the RMP and QMP data. Wilcoxon-rank sum test was used to compare the ratio between depleted- and enriched species. In negatively associated diseases such as Crohn's disease and ulcerative colitis, the majority of depleted species in the patients increased their effect size and statistical significance in the QMP compared to the RMP, while enriched species decreased the statistical significance. It was the opposite in the positively associated diseases such as colorectal cancer and slow transit constipation. These results indicate that RMP-based analyses underestimate or overestimate disease-associated microbial species due to differences in microbial load between cases and controls, while QMP-based analyses reduce these biases.
